# Non-genetic inheritance of induced defense morphologies across multiple unexposed generations of *Daphnia lumholtzi*

**DOI:** 10.1101/2025.09.28.679097

**Authors:** Shannon N. Snyder, Walker C. Meyer, Shanie L. Jorgenson, Ethan B. Contreras, Tea Bland, William A. Cresko

**Affiliations:** Institute of Ecology & Evolution, University of Oregon, Eugene, OR, 97403, USA; Department of Bioengineering, Knight Campus for Accelerating Scientific Impact, Eugene, OR, 97403, USA

**Keywords:** Transgenerational inheritance, Phenotypic plasticity, Transgenerational Plasticity, Non-genetic inheritance, Epigenetic Inheritance, Armor Morphology, Bayesian hierarchical modeling, Inducible defenses

## Abstract

**Teaser Text:** Can the ghosts of environments past shape organismal phenotypes of future generations? In an experiment with clonal *Daphnia lumholtzi*, we reveal that predator-induced changes in body shape in one generation can persist across several unexposed generations, even when the predator signal is long gone. Exposing only the first generation to predator cues, we tracked morphological shifts through five generations of genetically identical individuals. We found that these induced plastic phenotypes persisted to the F4 generation before fading in F5. This discovery supports the importance of non-genetic inheritance and underscores the powerful interplay between ancestral and current environments in shaping organismal form and function.

Environmental variation can induce phenotypic changes through adaptation of populations and acclimation of individuals. While genetic adaptation creates persistent change within a population via allele frequency shifts, reversible plastic phenotypes can be inherited through non-genetic mechanisms. However, most transgenerational plasticity studies examine generations where direct embryonic or germline exposure to environmental cues cannot be excluded. Empirical evidence for persistence definitively beyond the critical threshold for distinguishing true transgenerational plasticity from in utero exposure remains scarce, particularly for vertebrate predator-induced defenses.

We measured phenotypic effects of vertebrate predator exposure in the clonally reproducing water flea, *Daphnia lumholtzi*, isolating environmental effects from genetic variation. We exposed the F0 generation to fish conditioned media, then measured morphological defenses in definitively unexposed F3, F4, and F5 generations characterizing the temporal dynamics of non-genetic inheritance. Predator-induced morphologies persisted through F4 before receding in F5, demonstrating that non-genetic effects extend well beyond the embryonically exposed F1 and germ cell exposed F2 generations. This provides rare empirical evidence for transgenerational plasticity lasting through F4, without confounding genetic variation. We also examined ontogenetic patterns of somatic and defensive trait development, revealing trait-specific temporal dynamics in transgenerational effect expression and decay.

These results highlight how current and ancestral environments interact to determine phenotypic variation across generations and underscores the ecological significance of non-genetic inheritance in natural populations, particularly for understanding population responses to environmental change and predator reintroduction. Characterizing molecular mechanisms underlying transgenerational phenotypic plasticity remains critical for predicting persistence and ecological consequences of non-genetic inheritance.

## Introduction

Organisms respond to environmental variation via changes in morphology, behavior, and life history traits (Donelson et al., 2018; Szűcs et al., 2019; West-Eberhard, 2005). These phenotypic responses may arise from shifts in allele frequencies across generations or from non-genetic mechanisms acting within individuals (Levis & Pfennig, 2019; Uller et al., 2018; West-Eberhard, 2020). Increased phenotypic variation can enhance the probability that some offspring will express trait values suited to novel environmental conditions (Botero et al., 2015; Simons, 2009; Tufto, 2015). However, this variation also increases the likelihood of mismatched phenotypes, reducing individual fitness. While genetic adaptation can improve population-level fitness over time, subsequent environmental shifts may render previously adaptive traits maladaptive (Lande, 2009; Radchuk et al., 2019).

Regularly experienced environmental variation presents a particular challenge for evolving populations because it can lead to persistent mismatch of organisms and the environment, and thus ongoing genetic load (Ghalambor et al., 2007; Via et al., 1995). An alternative biological strategy for dealing with such variable conditions is the evolution of plastic traits (Hendry, 2016) in which phenotypic expression covaries with environmental variation (Bradshaw, 1965; Schlichting & Levin, 1986). Phenotypic plasticity allows a single genotype to consistently produce different phenotypes depending on the environment, without requiring genetic change (Pfennig et al., 2010; Pigliucci et al., 2006). Plasticity includes developmental plasticity (Nettle & Bateson, 2015; Uller et al., 2018), where early-life cues shape long-term traits, and acclimatory plasticity (Angilletta, 2009; Beaman et al., 2016), involving reversible physiological changes in response to short-term fluctuations, illustrating a range of temporal scales over which environmentally induced phenotypic changes can occur.

Plasticity within a generation has been well documented across numerous taxa and environmental conditions (Agrawal et al., 1999; Corl et al., 2018; Forsman, 2015). Although plasticity itself is often an adaptive phenotype that has been crafted by natural selection, the trait distributions produced by plasticity within one generation have historically been thought to be ephemeral with little consequence for the phenotypes of future generations (West-Eberhard, 2020). However, modern analyses have documented that variation in plastic traits is mediated by changes in gene expression (Hales et al., 2017) and is hypothesized to involve epigenetic regulation (St-Cyr & McGowan, 2015), raising the possibility that induced plastic phenotypes may be inherited across generations (Harris et al., 2012a; Romero-Mujalli et al., 2024). This potential for non-genetic inheritance exemplifies the need to integrate genetic and non-genetic mechanisms of information transmission across generations (inclusive heritability, Danchin & Wagner, 2010) in evolutionary biology.

Transgenerational inheritance of induced plastic traits from parent to offspring is expected to evolve when the parental environment covaries with the offspring environment (Agrawal et al., 1999). Although induced plastic phenotypes should return to the baseline state in the absence of the inducing cues over generations, the rate at which this occurs is hypothesized to be determined by the frequency of which the environment shifts (Furrow & Feldman, 2014; Proulx & Teotónio, 2017). The evolution of inherited phenotypic plasticity therefore provides a mechanism for organisms and their offspring to respond adaptively to changing environments without significant genetic load (Ghalambor et al., 2007).

Studies have documented that induced phenotypes can be inherited (Bell & Hellmann, 2019; MacLeod et al., 2022; Moore et al., 2019; Neylan et al., 2025; Tariel et al., 2020; Yin et al., 2019), but the prevalence and duration of this phenomenon remain unclear, especially under fluctuating environmental conditions (Norouzitallab et al., 2014). A particular challenge is distinguishing true transgenerational inheritance from extended maternal or embryonic effects. Because embryos often exist within the maternal body or egg, they may be directly exposed to the environmental cue (Agrelius & Dudycha, 2025; Skinner, 2008). Furthermore, these embryos may have developing germ cells that can be similarly exposed, necessitating the examination of generations sufficiently removed from the initial exposure to rule out direct effects. The number of generations required to differentiate maternal effects from true transgenerational inheritance varies by organism depending on developmental timing and reproductive mode, but empirical data demonstrating persistence beyond the last generation with possible germline exposure remains limited, partially because embryonic and germ line exposure is often conflated with true transgenerational plasticity (Agrelius & Dudycha, 2025; Skinner, 2008). In fact, a 2020 meta-analysis of transgenerational phenotypic plasticity (TGP) in predator-prey interactions found that only 13% of studies (seven total studies) extended beyond the F0 and F1 generations (Tariel et al., 2020), with studies examining F3+ generations remaining notably rare. Despite these advances, critical questions remain: How long do induced traits persist without ongoing cue exposure? At what generation do transgenerational effects decay? And can we definitively distinguish transgenerational plasticity from extended maternal effects?

Convincing examples of non-genetic inheritance that extend beyond temporal limits of potential direct exposure include a handful of examinations in *Caenorhabditis elegans* (Greer et al., 2011; S. W. Kim et al., 2013; Kishimoto et al., 2017; Klosin et al., 2017; Remy, 2010; M. Wang et al., 2019), *Mus musculus* (Anway et al., 2005; Skinner et al., 2013; Van Steenwyk et al., 2018), *Drosophila melanogaster* (Panacek et al., 2011), *Danio rerio* (Alfonso et al., 2019), *Ovis aries* (Kizilaslan et al., 2025), and the keystone aquatic species, *Daphnia* (Shahmohamadloo et al., 2025).

Often overlooked in such studies is the influence of genetic variation. Asexual reproduction provides a particularly powerful system for studying non-genetic transgenerational effects because it eliminates confounding variation due to recombination, genetic background, and parental sex. Examples of such studies include a striking example in aphids, which examined 27 generations of the inheritance of predation induced phenotypes (Sentis et al., 2018), clonal and hermaphroditic freshwater snail systems (Dejeux et al., 2025; Smithson et al., 2020; Tariel et al., 2020), in which alterations in shells shape and thickness can be inherited across generations, and hermaphroditic *C.elegans* (Baugh & Day, 2020). These examples remain relatively few, leaving it unclear whether true transgenerational inheritance is rare or understudied.

The freshwater microcrustacean *Daphnia* is especially well suited for studying transgenerational plasticity and addressing these questions (Ebert, 2022; Harris et al., 2012; Seda & Petrusek, 2011; Yampolsky et al., 2014). The concept of plasticity was introduced in the early 1900s by Richard Woltereck, who demonstrated that head size in *Daphnia* varied with nutrient availability, and proposed that environmentally induced traits could be inherited (Woltereck, 1913). *Daphnia’s* parthenogenetic reproduction produces genetically identical offspring, eliminating confounding variation due to recombination, genetic background, and parental sex (Harris et al., 2012). *These ecoresponsive crustaceans* exhibit plastic responses to a range of environmental stressors, including cyanobacteria (Dao et al., 2018; Gustafsson & Hansson, 2004; Schwarzenberger & Martin-Creuzburg, 2021), temperature (Geerts et al., 2015; Landy et al., 2024; Müller et al., 2018), and predation (Hebert & Grewe, 1985; Tollrian & Harvell, 1999), affecting morphology (Horstmann et al., 2021), behavior (Beklioglu et al., 2006), and life history traits (Adamczuk, 2020; Agrawal et al., 1999). Examples include inducible helmets, spines, and neckteeth in response to predator cues (Horstmann et al., 2021; Laforsch & Tollrian, 2004). Many of these traits persist across clonal generations, such as helmet length in *D. cucullata* (Agrawal et al., 1999) and clutch size in *D. ambigua* (Walsh et al., 2014), suggesting a non-genetic mechanism of inheritance.

The number of generations over which induced traits persist in *Daphnia* in the absence of the original environmental cue is still being resolved. While phenotypic effects extending through the F1-F2 generations (maternal effects) in *Daphnia* are well-documented, evidence for persistence beyond F2 (which eliminates direct embryonic exposure; (Agrelius & Dudycha, 2025) remains limited. Most documented cases track only one or two generations post-exposure, such as helmet length in *D. cucullata* (Agrawal et al., 1999; Laforsch & Tollrian, 2004), and age of maturation in *D. ambigua* (Walsh et al., 2016). Many of these studies have predominantly focused on invertebrate predation, such as *Chaoborus* larvae (Horstmann et al., 2022), *Bythotrephes longimanus* (spiny water flea) (Landy et al., 2024), *Leptodora* (Baludo et al., 2024), *Notonecta* (Horstmann et al., 2023; Weiss et al., 2015), and *Triops* (Pietrzak et al., 2020), or focus on various toxicological stressors (Funke et al., 2024; J. Kim & Choi, 2023; Liao et al., 2023; Liu et al., 2020; Maggio & Jenkins, 2022a, 2022b; Martins & Guilhermino, 2018a, 2018b; Shahmohamadloo et al., 2025; Song et al., 2022; Vandegehuchte, Lemière, et al., 2010; J. Wang et al., 2024). However, studies examining the transgenerational responses to vertebrate predation beyond generations with possible germline exposure remain notably scarce (Tariel et al., 2020).

This gap is particularly important because vertebrate predators impose strong, and often more temporally consistent selective pressure on prey populations (Batabyal, 2023; Salo et al., 2010; Wooster, 1994), making transgenerational responses to fish predation critical for understanding adaptive potential in natural systems, and relevant to predator reintroduction efforts (Bracis & Wirsing, 2021). Molecular studies suggest long term effects are possible, with differential gene expression persisting after predator cue removal (Hales et al., 2017), and methylation differences between naive and unexposed individuals across various stressors (Feiner et al., 2022; Hearn et al., 2019; Jeremias et al., 2018; Trijau et al., 2018). However, the duration of phenotypic persistence and its evolutionary significance remain unclear (Bird, 2024; Feinberg, 2007; Feinberg & Irizarry, 2010; Fitz-James & Cavalli, 2022; Ho & Burggren, 2010; Klibaner-Schiff et al., 2024), requiring empirical evidence beyond generations with possible germline exposure.

*Daphnia lumholtzi* develops elongated head and tailspine in response to fish predation (Dzialowski et al., 2003), traits that effectively reduce predation risk and represent a clear example of adaptive phenotypic plasticity (Engel et al., 2014). These distinctive spines are thought to contribute to the species’ invasive success in North America (Engel & Tollrian, 2009; Sousa et al., 2017; Work & Gophen, 1999) where *D. lumholtzi* often displaces native North American *Daphnia* populations (Johnson & Havel, 2001). Unlike *D. lumholtzi,* many North American *Daphnia* species typically produce broad, helmet-like adornments rather than elongated spines (Horstmann et al., 2021) exemplifying *D. lumholtzi’s* unique defensive strategy. These distinctive morphological defenses, combined with *Daphnia’s* parthenogenetic reproduction, make *D. lumholtzi* an ideal system for tracking transgenerational effects across multiple generations without genetic confounds.

We measured predator induced morphological traits in *D. lumholtzi* across multiple genetically identical generations following initial exposure, focusing on F3-F5 cohorts where neither the individuals themselves nor their germ cells (as developing gametes in F1-F2) could have experienced direct exposure to predator cues. Previous work in *D. lumholtzi* shows that continued multigenerational maternal exposure can accelerate defensive trait development (Graeve et al., 2021) and that cyanobacteria-induced life history effects persist for at least one generation (Dao et al., 2018), but no study has examined the long-term persistence of vertebrate predator induced morphology in *Daphnia lumholtzi*, particularly beyond the generations with possible direct exposure. By focusing on F3-F5 generations, our observations document transgenerational inheritance via necessarily non-genetic mechanisms rather than extended effects of direct exposure. Additionally, deriving all experimental lineages from a single founding mother of the Saguaro Lake *D. lumholtzi* clone eliminates confounding effects of standing genetic variation, enabling us to fully isolate non-genetic inheritance.

We found that induced traits persisted through multiple unexposed generations, with measurable effects gradually declining to undetectable levels in F5. We identify distinct ontogenetic trajectories in defensive versus somatic traits across the Daphnia’s lifespan. A critical question is whether transgenerational morphological difference reflect inherited plasticity versus size-selective mortality of non-induced phenotypes. To address this alternative hypothesis, we explicitly test for evidence of trait-selective or size-selective mortality and report overlapping survival trajectories between different sized defensive structures, indicating that elongated spines are indeed inherited, rather than a selective artifact. These results demonstrate that true transgenerational plasticity can be inherited across clonal generations, but exhibits limited temporal persistence, with trait-specific patterns of decline that reflect the gradual loss of ancestral environmental influence (Geoghegan & Spencer, 2012; Heard & Martienssen, 2014).

## Methods

### Experimental Overview (Figure 1)

To explore the dynamics of predation induced phenotypes within and across generations, we reared individual Daphnia with ancestral predation exposure in the F0 generation, alongside unexposed *Daphnia lumholtzi* individuals for five generations following initial predation exposure only in the predator F0 generation (total generations: F0-F5; only F0 exposed). These individuals were the progeny of a single founding mother, whose genetically identical daughters (F0) were randomly assigned to treatment or control to establish replicates, and propagated for an additional four generations without subsequent exposure. This design isolated environmental effects from confounding genetic variation while maintaining independent biological replicates.

**Figure 1.**
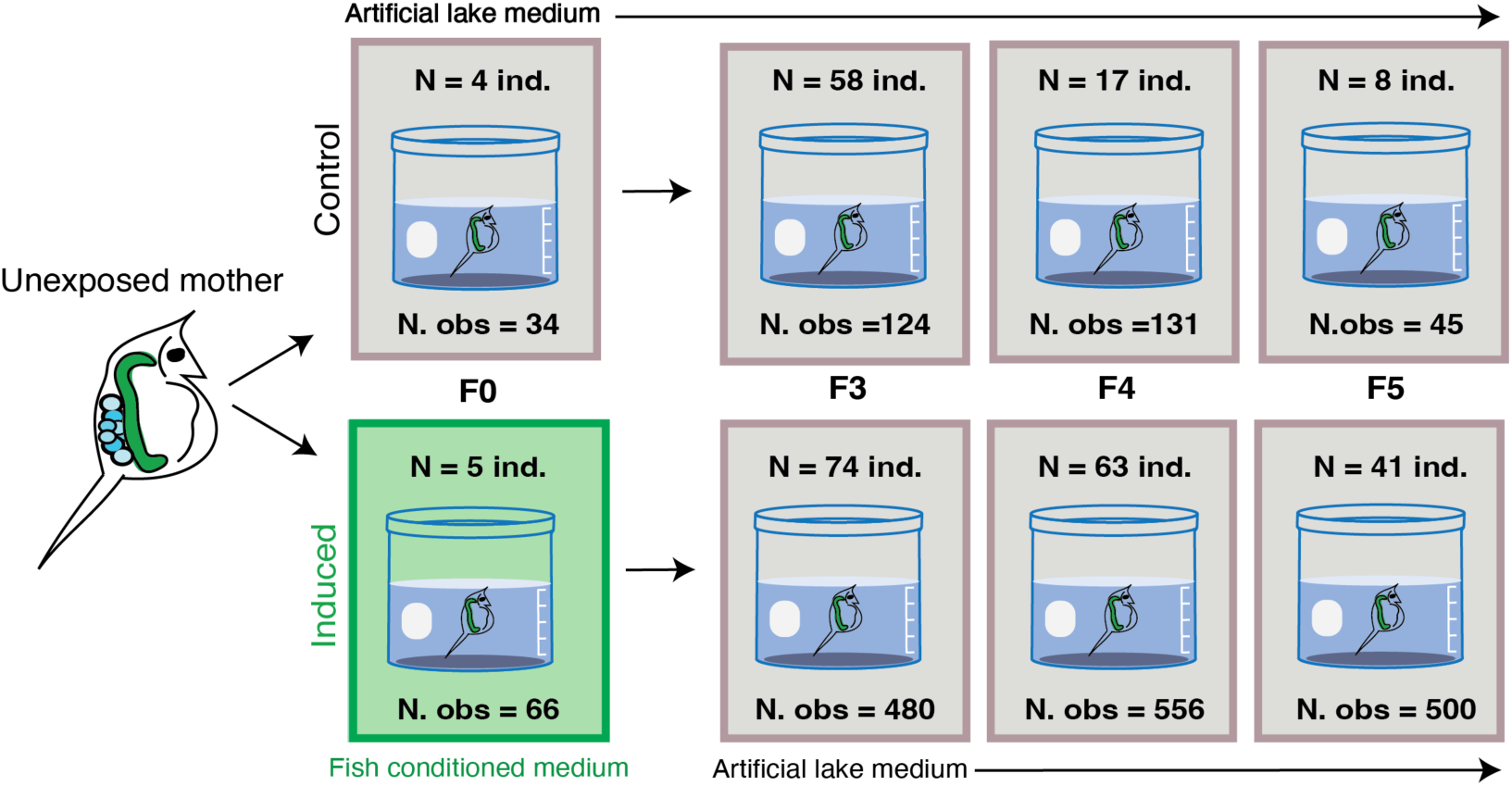
Schematic illustrating the experimental setup allowing for direct testing of transgenerational epigenetic inheritance by induced defenses by comparing individuals with and without grandparental exposure to predator cues. Offspring (F0) from a single unexposed mother were split into two genetically identical groups: control (top), where individuals where continuously reared in artificial lake medium, and induced (bottom), where F0 individuals were exposed to fish-conditioned medium indicated in green. Only the F0 induced generation is exposed to fish conditioned media. Sample size for each generation and treatment is indicated (N ind.), and the number of observations (repeated measures, N.obs) is indicated. Subsequent generations (F3-F5) from both treatment group and control are maintained in artificial lake media. Generations were propagated for six generations using the first viable (successful reproduction) and measured every other day during the F0, F3, F4, and F5 generations.

We focused morphological measurements on F0, F3, F4, and F5 generations. F0 established baseline induction effects, while F3-F5 represented generations with no direct embryonic or germline exposure to predator cues - essential for distinguishing true transgenerational inheritance from extended maternal effects (Agrelius & Dudycha, 2025; Skinner, 2008). F1 and F2 generations were propagated but not measured, as phenotypic effects in these cohorts are well-documented and can result from direct embryonic exposure. While this design limits F0 sample size, our primary objective was characterizing persistence of effects in F3-F5 individuals rather than recapitulating characterized F0-F2 responses.

*Daphnia* were individually grown in 50 mL beakers containing 30 mL of fish conditioned medium or control medium (COMBO) (Kilham et al., 1998), enabling replication at the individual level. Every other day, until death, during the F0, F3, F4 and F5 generations, individual *D. lumholtzi* were imaged for morphological measurements using `ImageJ` (Schindelin et al., 2012). This experimental framework also allowed us to examine survivorship (as animals were imaged until they died) and life-long ontogenetic trajectories. On this same schedule the media was exchanged, and algae was replenished with 85,000 cells/mL of a 1:1 mixture of *Scenedesmus obliquus* and *Chlorella vulgaris*, hereafter referred to as the standard algae mixture.

Variation in sample size between treated and control individuals and across generations reflects differential survival and reproductive success. Predator-exposed individuals showed higher survival and fecundity in some generations, resulting in asymmetric sample sizes. Bayesian hierarchical models can maintain robustness despite imbalance and small sample sizes. See Figure 1 for experimental schematic and sample sizes; See Table 1 for expanded sample sizes, including number of repeated measures observations on individuals.

**Table 1:** Sample sizes across experimental generations and treatment showing number of individuals and repeated observation details.

| Generation | Treatment | Individuals | Observations | Mean Obs/Ind |
| --- | --- | --- | --- | --- |
| F0 | induced | 5 | 66 | 13.2 |
|  | control | 4 | 34 | 8.5 |
| F3 | induced | 74 | 480 | 6.5 |
|  | control | 58 | 124 | 2.1 |
| F4 | induced | 63 | 556 | 8.8 |
|  | control | 17 | 131 | 7.7 |
| F5 | induced | 41 | 500 | 12.2 |
|  | control | 8 | 45 | 5.6 |

This study focuses on a single genetic background to isolate non-genetic effects. Future work examining multiple clones and populations will be necessary to assess the generalizability of these transgenerational effects.

### Clonal Genetic Background: *Daphnia Lumholtzi*

*Daphnia lumholtzi* was collected in the summer of 2019 from Saguaro Lake, AZ, USA. We utilized one Saguaro Lake Daphnia, as our primary objective is to minimize confounding genetic variation, rather than generalize a population’s ability to respond to predator cues. This species and genetic background were specifically chosen for its ability to develop unique headspines and tailspines in response to vertebrate predation and for their ecological significance as an invasive North American species, capable of displacing native *Daphnia* species (Engel et al., 2014b; Engel & Tollrian, 2009; Sousa et al., 2017; Work & Gophen, 1999).

### Common Garden Rearing to Standardize Generational Exposures

After lake collection, *D. lumholtzi* were reared for 6 months in artificial lake medium (COMBO) and fed *Scenedesmus obliquus* and *Chlorella vulgaris* ad libitum to dull residual effects of lake predation. Prior to experimental perturbations, *D. lumholtzi* were common garden reared for three generations in COMBO medium (Kilham et al., 1998) and fed a standard algae mixture of *S. obliquus* and *C. vulgaris* to minimize the influence of maternal effects on experimental generations.

### Fish and Control Media Preparation

To prepare fish (induction) medium, 10L of artificial lake medium (COMBO) was conditioned with five *Gasterosteus aculeatus* for three days prior to filtration (0.45 μm). Fish were fed 10 live *D. lumholtzi* and 10 *D. magna* daily to provide both conspecific and heterospecific prey cues, maximizing predation kairomone concentration. To create control medium, artificial lake medium without fish conditioning was created in parallel with fish medium. Media was frozen and thawed overnight prior to use.

### Initial F0 Predation Exposure and Experimental Conditions

From the third generation of common garden reared *Daphnia*, one mother was selected due to observed clutch survivability and clutch size. Nine neonates born within a 12 hour period from the same mother were isolated and randomly assigned to treatment groups, either fish exposure or control. Animals were reared individually in 50 mL plastic beakers with 30mL of the appropriate medium and fed 5 mL standard algae mixture every other day, ensuring replication at the individual level. Individual Daphnia were staged, imaged, and returned to fresh media with algae every other day for the duration of their lives.

### Propagation across Generations

Upon reproduction, up to six neonates from each mother’s first clutch were randomly collected and reared individually in 50 mL beakers containing 30 mL control medium. If individuals from the mother’s first clutch did not survive to reproduce, neonates from the mother’s subsequent clutch were used instead. This ensured propagation of all lineages while maintaining consistency in clutch selection across treatments. See Figure 1 and Table 1 for samples sizes.

### Imaging and Data Collection

Every other day, animals in the F0, F3, F4, and F5 generation were staged on a glass slide and imaged using `cellSenseEntry` (Olympus Life Science). Scale bars were added to images during acquisition for calibration during measurements.

Morphological measurements of *headspine* (tip of spine to top of compound eye), *body length* (top of eye to base of tail), *tailspine* (base of tail to tip of tailspine), *total body length* (tip of headspine to tip of tailspine) were collected (see Figure 2) using `ImageJ` (Schindelin et al., 2012). All images were calibrated to a consistent scale using the ‘set global scale’ function in `ImageJ`, which converts pixel distances to micrometers based on the embedded scale bar. See Figure 2 for measurement details.

**Figure 2.**
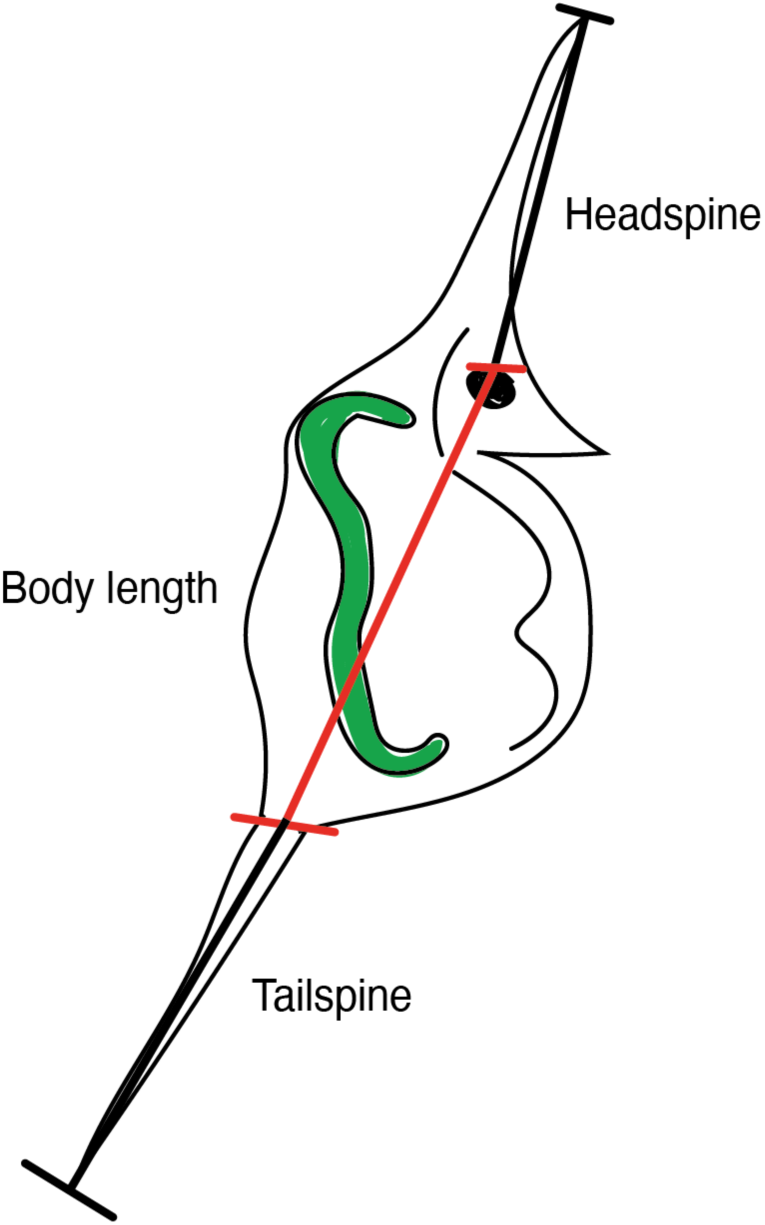
Measurement of inducible morphological defenses in Daphnia in response to predator cues. Daphnia reared in fish-conditioned media develop significantly elongated headspines and tailspines compared to control animals. Representative lateral images of morphological measurements are indicated, detailed as follows: Headspine was measured from tip of spine to top of compound eye, body length was measured from top of compound eye to base of tailspine, tailspine was measured from base to tip of spine. Total body size was measured from top of headspine to tip of tailspine.

### Ontogenetic Trajectories

To illustrate general patterns of ontogenetic development in somatic and defensive traits, Figure 3 presents scatterplots pooled across generations, using `ggplot2` (Wickham, 2016). Generation- and treatment-specific effects are subsequently analyzed in detail in our primary statistical models (see Primary Hypothesis section below).

**Figure 3.**
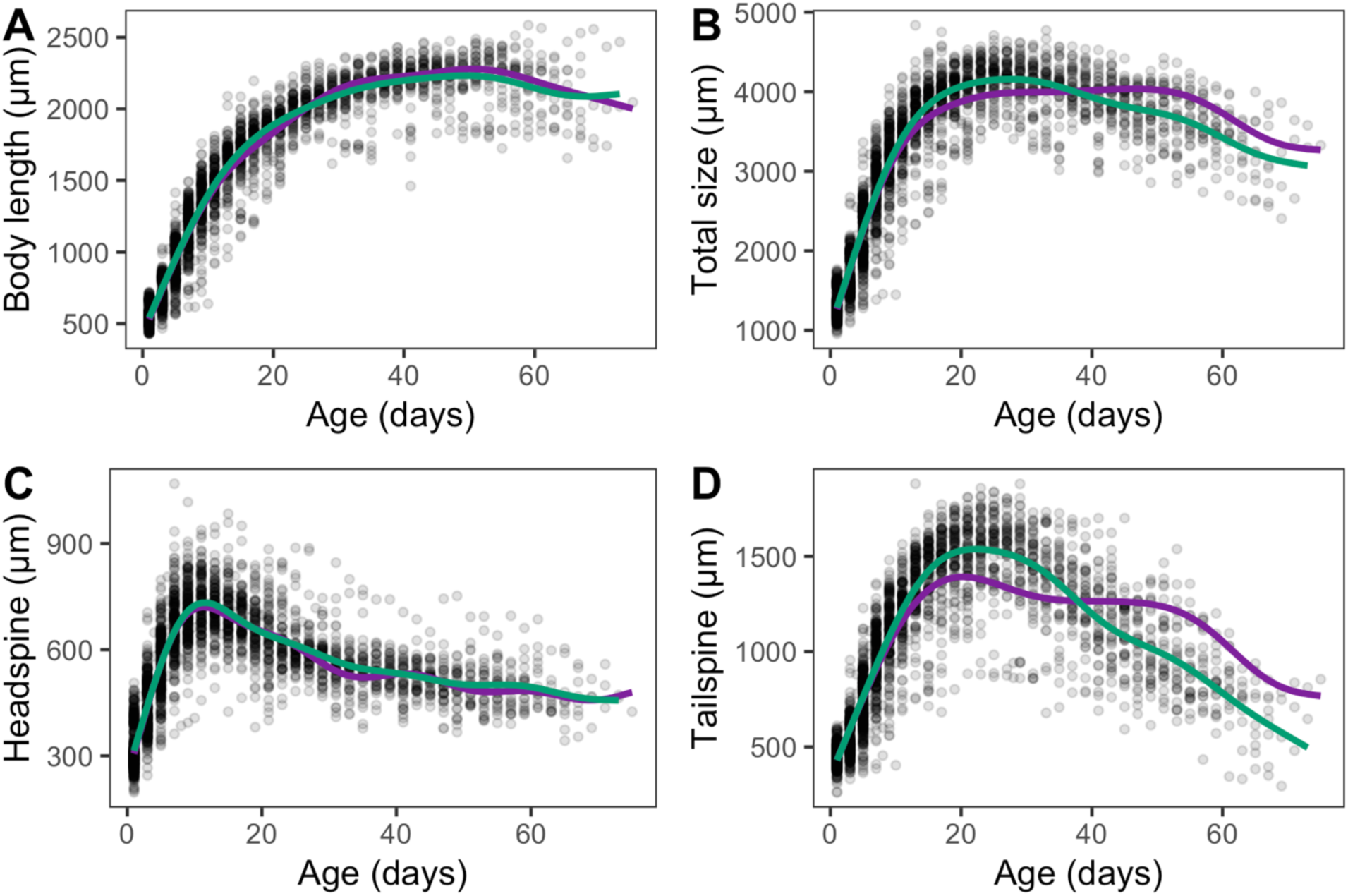
Trajectories of somatic and defensive traits reveal divergence in late-life dynamics. Scatterplots display individual measurements of (A) body length, (B) total body size, (C) headspine length, and (D) tailspine length (in μm) as a function of age (in days). Green line represents locally averaged (LOESS) trends for induced individuals (F0 exposed to fish-conditioned media); purple line represents locally averages (LOESS) trends for control individuals (no F0 exposure). Somatic traits - body length (A) and total body size (B) - increase rapidly during early juvenile stages, then reach a plateau in mid-life, with little elongation even at advanced ages. In contrast, defensive traits - headspine (C) and tailspine (D) – regress later in life. Morphological trajectories demonstrate distinct patterns in defense trait expression, but not body growth, between control and treated individuals, consistent with the inheritance of plastic response initiated by ancestral exposure.

### Statistical Analysis

#### Primary Hypothesis: Phenotype Transmission Across Generations

Because individuals were measured repeatedly within generations, we used hierarchical mixed models to account for non-independence. We employed complementary Bayesian hierarchical and frequentist mixed-effects approaches to ensure robust inference. Bayesian models provide full posterior distributions enabling probabilistic statements about treatment effects and improved estimation with sparse data through partial pooling (Gelman & Hill, 2006). Frequentist models provide p-value-based hypothesis testing for readers familiar with traditional frameworks. Importantly, both approaches yielded consistent effect size estimates and overlapping confidence/credible intervals across all generations and traits (Table 2, Supplemental Table 2), supporting the reliability of our conclusions regardless of statistical framework.

**Table 2.** Posterior mean estimates of treatment-induced changes in spine length across generations.

| Spine | Generation | Estimate | SE | 95% CrI | PD | R<sup>2</sup> (marg.) |
| --- | --- | --- | --- | --- | --- | --- |
| Head Spine | F0 | 68.54 | 32.90 | [1.67, 132.9] | **97.7%** | 60.4% |
|  | F3 | 38.19 | 18.32 | [3.1, 74.71] | **98.5%** | 64.9% |
|  | F4 | 44.36 | 20.45 | [3.99, 84.58] | **98.4%** | 65.8% |
|  | F5 | -7.07 | 22.32 | [-51.23, 36.78] | 62.6% | 56.5% |
| Tail Spine | F0 | 99.71 | 41.28 | [15.35, 181.54] | **98.6%** | 94.5% |
|  | F3 | 5.81 | 18.35 | [-30.24, 41.81] | 62.5% | 98% |
|  | F4 | 85.57 | 34.76 | [16.95, 154.28] | **99.3%** | 98.3% |
|  | F5 | -19.21 | 29.07 | [-77.17, 37.5] | 74.7% | 94.8% |

Models were fit separately by generation for individuals <20 days old. Each row represents a unique combination of generation and spine type. 'Estimate' is the posterior mean difference in spine length ( $\mu\text{m}$ ) between predator-induced and control treatments, with positive values indicating elongation in induced individuals. 'SE' is the posterior standard deviation. '95% CrI' denotes the 95% credible interval for the posterior distribution. 'PD' (Probability of Direction) represents the posterior probability that the effect is in the estimated direction (positive indicating higher trait values in treated, and negative indicating lower trait values in treated, compared to control). 'R<sup>2</sup> (marg.)' is the marginal Bayesian R<sup>2</sup>, indicating variance explained by fixed effects only. See Methods for full model specification and prior structure.

Morphological variation was analyzed using both frequentist and Bayesian mixed-effects models in `R` (R Core Team, 2021), using the `lme4` (Bates et al., 2015) and `brms` (P. C. Bürkner, 2017) packages respectively. For each generation (*F0, F3, F4, F5*) and each trait (*headspine, tailspine*), we tested for the effects of predation treatment, with *body length* as a covariate, and a random intercept for individual identity to account for repeated measures on individuals. The model formula used was:

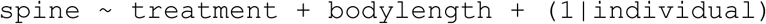

where *individual* represents the individual Daphnia identity, accounting for repeated measures on the same organism over time. The *bodylength* term is a covariate accounting for differences in body length that contribute to spine length, and treatment represents whether the F0 progenitor was exposed to predation (for F3, F4 and F5); F0 treated animals were directly exposed. Spine refers to either the measured head or tailspine length.

We selected this model structure based on established principles in *Daphnia* morphological analysis and our experimental design. *Body length* was included as a covariate because spine length scales allometrically with body size in *Daphnia* and failing to account for this relationship can confound treatment effects with size-related variation (Laforsch & Tollrian, 2004; Petrusek et al., 2009). *Treatment* (predator exposure vs. control) was included as the main fixed effect to test our primary hypothesis of transgenerational inheritance of induced defenses. The random intercept for individual identity *(1|individual)* accounts for repeated measurements on the same individuals, addressing potential pseudo-replication issues common in experimental ecology (Freeberg & Lucas, 2009; Hurlbert, 1984). We included *body length* rather than *age* as a covariate because they are highly correlated (Pearson’s r = 0.85; Supplemental Figure 1), and *body length* more directly captures the allometric relationship between body size and spine development (Laforsch & Tollrian, 2004).

#### Bayesian Hierarchical Models

We implemented Bayesian hierarchical models with the model structure described above. The hierarchical structure allows partial pooling across individuals, improving estimation in the presence of sparse data. This method also improves upon *p*-value based frequentist methods by providing a full posterior distribution for the effect size, allowing probabilistic statements about parameter values rather than relying on a binary significant/non-significant framework. Models were fit using a Student’s t-distribution to account for potential outliers and heavy-tailed data, which are common in ecological trait data. We used weakly informative priors and ran three chains with 5,000 iterations each (2,000 warm-up), ensuring convergence (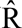 < 1.01 for all parameters). Posterior distributions were summarized to extract treatment effect estimates and 95% credible intervals. We generated posterior predictions at the mean body length using `posterior_epred()`from the `posterior` package (Bürkner et al., 2025), isolating treatment effects while controlling for individual-level variation. These predictions are visualized in Figure 4. We determined adequate model fit and convergence using 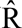 values, effective sample sizes (ESS), leave-one-out cross-validation (LOOIC), WAIC (Supplemental Table 1; (Vehtari et al., 2021), trace plots, and posterior predictive checks (Supplemental Figure 2 and 3; Gabry et al., 2019). Marginal and conditional R² values were computed using the `performance` package (Lüdecke et al., 2021) to assess model fit. Treatment effect estimates, including standard errors, confidence intervals, probability of direction (PD), and R² values are presented in Table 2. The predicted effect of treatment on defensive traits across generations is visualized in Figure 4.

**Figure 4.**
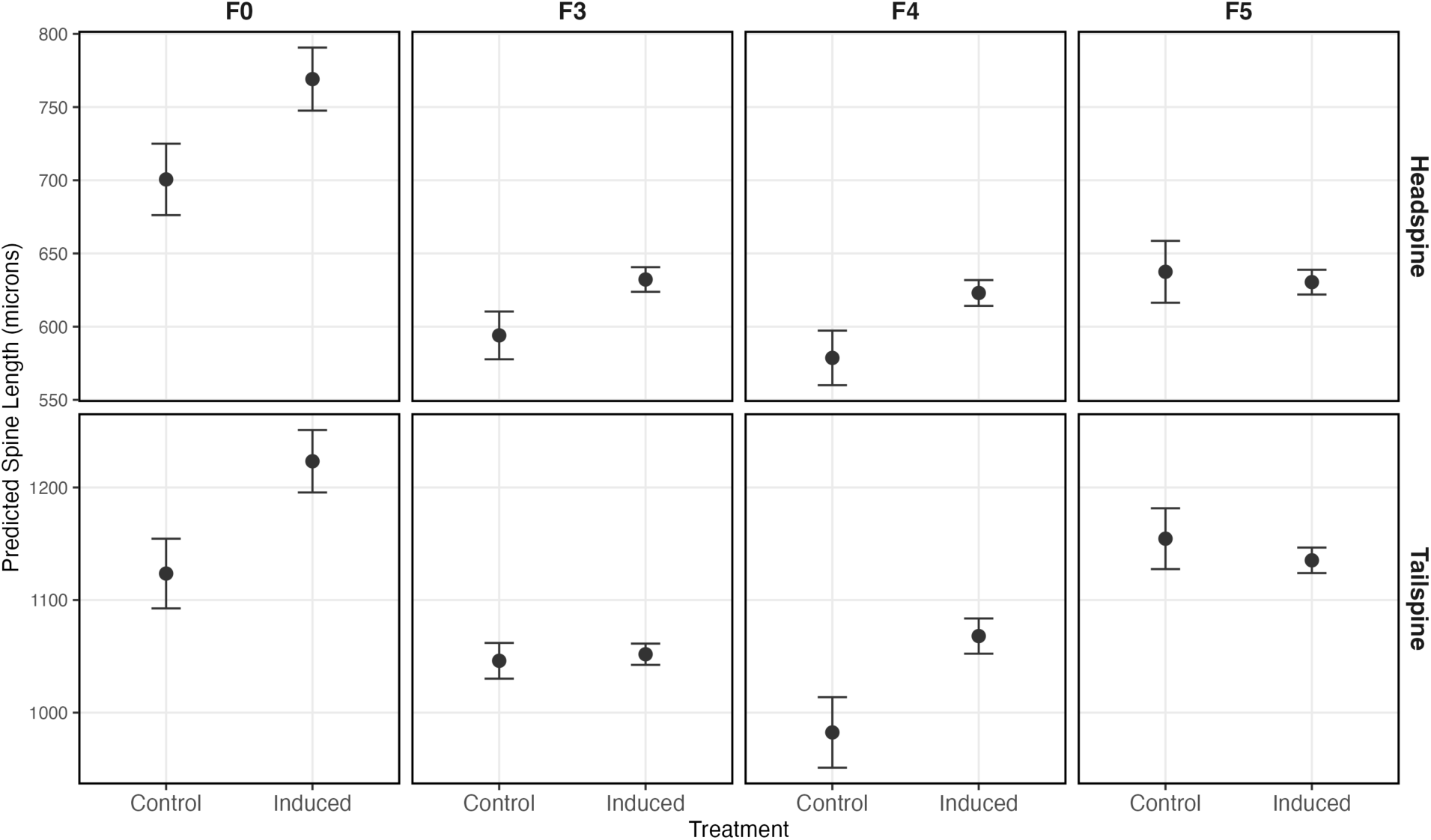
Predicted spine lengths by generation and treatment. Predicted mean headspine length (top row) and tailspine length (bottom row) for *Daphnia* in control and induced individuals across generations (F0, F3, F4, F5). Points represent model-predicted means ± standard error. Estimates are based on data from individuals ≤ 20 days old when defensive traits are most pronounced (Figure 3). See Table 2 for underlying estimates and parameters, and Supplemental Table 1 for model diagnostics.

#### Frequentist Mixed-Effects Models

To provide familiar p-value-based hypothesis testing, we also conducted frequentist analysis using the `lme4` (Bates et al., 2015) `R` package with the same model formula. Linear mixed models were evaluated using Kenward-Roger F-tests and post hoc Tukey tests to assess treatment effects. Conditional and marginal R² values, AIC, and BIC were calculated to assess model fit (Supplemental Table 3). Bayesian and frequentist estimates were highly consistent across generations and traits, with overlapping intervals and similar effect sizes (Table 2; Supplemental Table 2). For transparency, raw means comparisons are presented in Supplemental Figure 5.

#### Primary Analysis: Testing for Size-Dependent Treatment Effects

To specifically assess the effect of treatment on *body length* across generations, we again utilized Bayesian hierarchical modeling using the following formula:

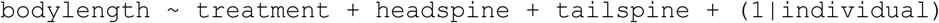

This model determines whether *treatment* affects body size independent of spine elongation. We used *spine* lengths (and not *age*) as covariates because *age* and *body length* are highly correlated (Pearson’s r =0.85, Supplemental Figure 1) and because our spine models used *body length* (not *age*) as a covariate, making this specification more consistent. As with the spine models, we used weakly informative priors and a Student’s t distribution and ran 2000 warmup and 4000 iterations using 4 chains and 4 cores utilizing the `brms R` package (Bürkner, 2017). Trace plots and posterior predictive checks confirmed model convergence and adequate model fit and are included in the supplemental materials for reference (Supplemental Figure 6).

#### Secondary Analysis: Age-Dependent Morphological Plasticity (Delta)

To assess how morphological traits change across *age* and *treatment*, we calculated within-individual morphological change (Δ) for *body length*, *total body size*, *headspine*, and *tailspine*. Δ values were computed as the difference between successive measurements for each individual (`Δtrait = trait_current − trait_previous`), capturing short-term plastic responses over time. Age trajectories (*Δ trait* vs. *age*) were visualized using scatterplots with LOESS-smoothed trend lines and 95% confidence intervals (Figure 5). To statistically evaluate the effect of induction on morphological plasticity, we modeled Δ values using linear mixed-effects models, `lme4` (Bates et al., 2015) and `lmerTest` package (Kuznetsova et al., 2017), with *treatment* and *age* as fixed effects and individual identity as a random intercept to account for repeated measures:

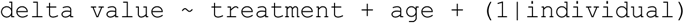

**Figure 5.**
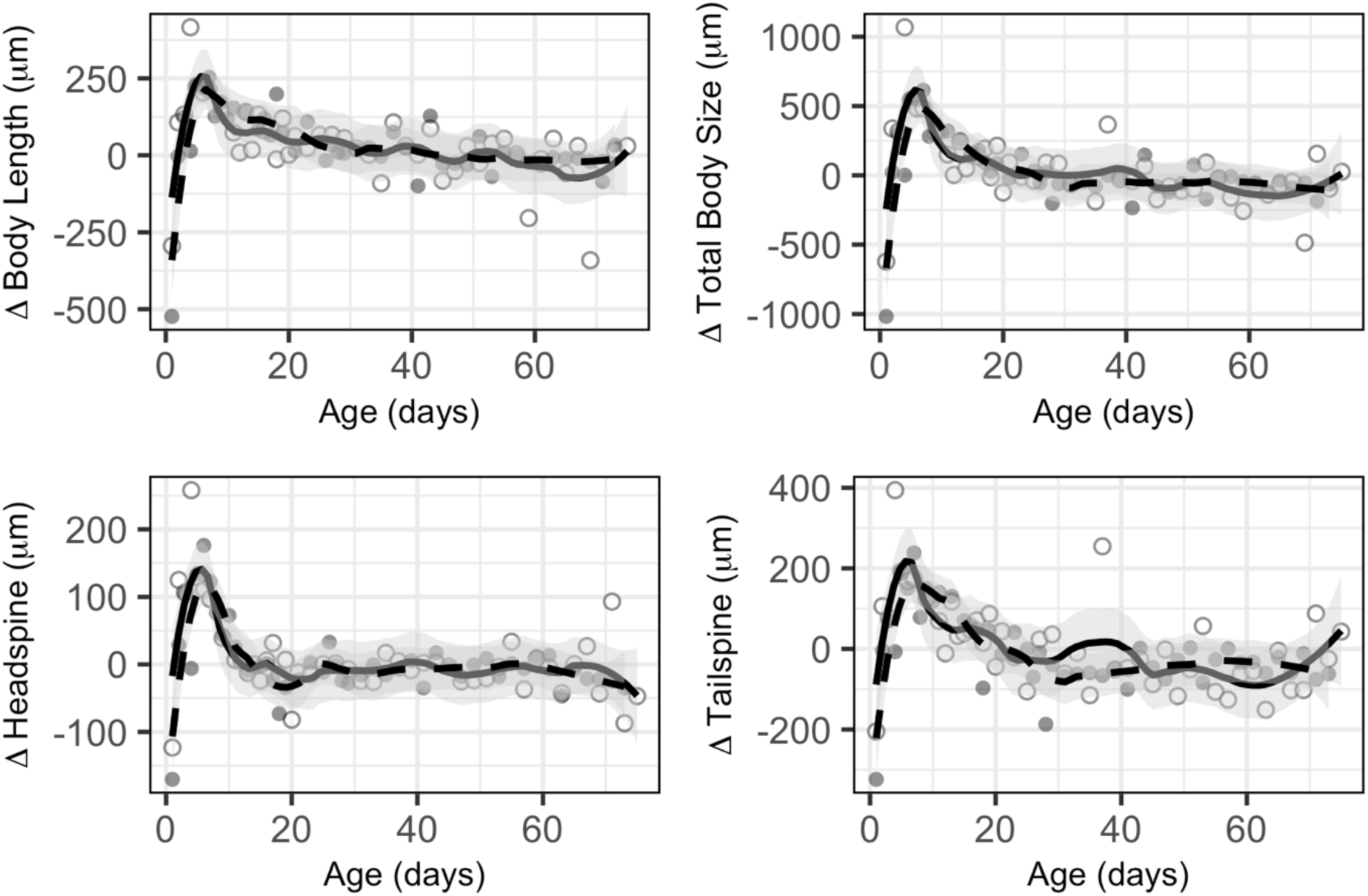
Growth of defensive and somatic traits are most pronounced early in life, followed by a stabilization with age, and were unaffected by treatment. Changes in morphological measurements (body length, total body size, headspine, and tailspine) across the lifespan (days). Each panel shows the mean growth (Δ; in μm) from the previous timepoint for the indicated trait. Morphological changes are largest early in life and stabilize as individuals age, as shown by the bold smoothed trend line with 95% confidence intervals (shaded area). Induced/treated animals are represented by the solid line, and control animals are indicated by the dashed line.

Because this analysis addressed a secondary question about short-term morphological plasticity rather than our primary hypothesis on spine inheritance, we opted for a parsimonious linear mixed-effects approach. This strategy ensures computational efficiency for secondary analyses while providing the level of inferential detail appropriate for each research question. We initially explored more complex temporal modeling approaches, including natural splines with varying degrees of freedom (df = 2 to 4) to capture potential non-linear age-related changes in growth patterns:

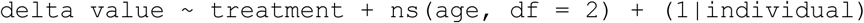

However, these models suffered from overparameterization, and model significance was sensitive to the choice of degrees of freedom, indicating unstable parameter estimates. Consequently, we adopted the more parsimonious linear approach with *age* as a continuous covariate, which provided robust and interpretable results while adequately capturing the age-related trends in our data. Results were visualized in Table 3, including the estimated effect of treatment, standard error, F-statistic, and *p-*value. For a more granular approach, we also modeled these effects by generation and present the same parameters (Supplemental Table 4).

**Table 3.** Treatment effects on body length controlling for spine morphology.

| Generation | Est. Difference (μm) | 95% CrI | Cohen's d | PD |
| --- | --- | --- | --- | --- |
| F0 | -0.53 | [-99.45, 95.21] | < 0.01 | 50.3% |
| F3 | -2.53 | [-39.42, 34.84] | -0.05 | 55.1% |
| F4 | -61.45 | [-122.54, 2.22] | -1.16 | 97.1% |
| F5 | 0.46 | [-93.78, 89.51] | 0.03 | 51.7% |

Treatment effects on body length across generations. Bayesian hierarchical models tested whether ancestral predator exposure affected body size independent of spine elongation, controlling for headspine and tailspine length, and with a random intercept for individual identity. Negative estimates indicate smaller body size in predator-exposed individuals. See Methods for full model specification. 'Generation' refers to the generation examined, 'Est. Difference (μm)' is the estimated effect of treated – control, therefore negative values represent a smaller estimated body size in treated animals. '95% CrI' depicts the 95% credible intervals. For interpretability, we include the estimated effect sizes in the 'Cohen's d' column. 'PD' (Probability of Direction) represents the posterior probability that the treatment effect is in the estimated direction. Values >97.5% indicate strong evidence for the effect direction; values near 50% indicate high uncertainty.

#### Secondary Analysis: Size Selective Survivorship Analysis

To test whether size-selective mortality, rather than treatment, could explain differences in spine lengths, we constructed mixed-effects linear models using binary survival status. Because this analysis addressed a secondary question about size-selective mortality rather than our primary hypothesis on spine inheritance, we opted for a parsimonious frequentist mixed-effects approach. Bayesian modeling was unnecessary here because the goal was to test a simple association between body size and spine length, and linear mixed-effects models provided interpretable estimates while accounting for repeated measures. This approach balances computational efficiency with the depth of inference appropriate for each research question. *Survival status* was defined as the presence or absence of a subsequent timepoint for each individual. Separate models were fit for *headspine* and *tailspine* lengths as a function of *body length*, with a random intercept for individual identity.

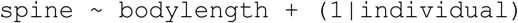

Residuals from these models represent spine length variation independent of body size. To visualize trait dynamics across age and survival status, we plotted body length and spine residuals using LOESS-smoothed trend lines with 95% confidence intervals. Survivors and non-survivors were distinguished by color, and traits were faceted for comparison (Figure 6).

**Figure 6.**
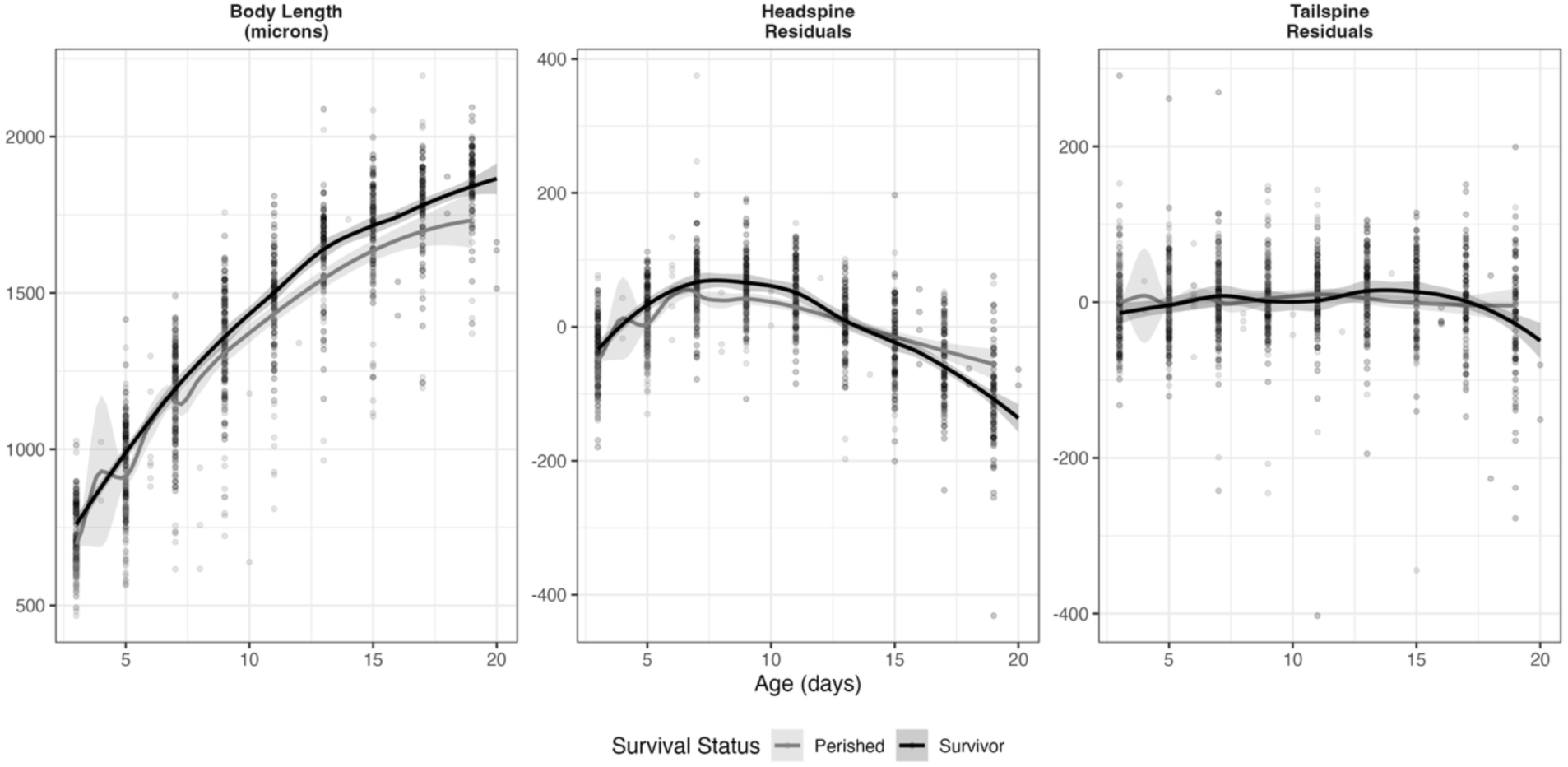
Developmental trajectories during periods of concentrated growth do not indicate treatment or size selective survivorship. Body length, head spine (residual), and tail spine (residual) measurements over first 20 days of age for individuals grouped by survival status (dark grey for those who survived to the next timepoint - survivors, light grey for those that perished). Residual values for head and tail spines are calculated from linear mixed-effects models that account for body length, isolating variation in spine length independent of body size. Both survivors and non-survivors display overlapping developmental trajectories and trait distributions, providing no evidence that size-selective or trait-selective mortality underlies observed group differences.

To further evaluate survival probability, we constructed a binomial generalized additive model (GAM) with *survival status* as the response variable and *age*, *body length*, *spine* lengths, treatment, and the interaction between *age* and *treatment* as predictors.

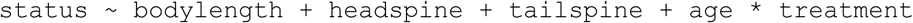

Trait values representing the 10th, 50th, and 90th percentiles were used to generate survival predictions across *age*. Predicted survival probabilities and 95% confidence intervals were plotted to assess how trait variation influences survivorship over time (Figure 7).

**Figure 7.**
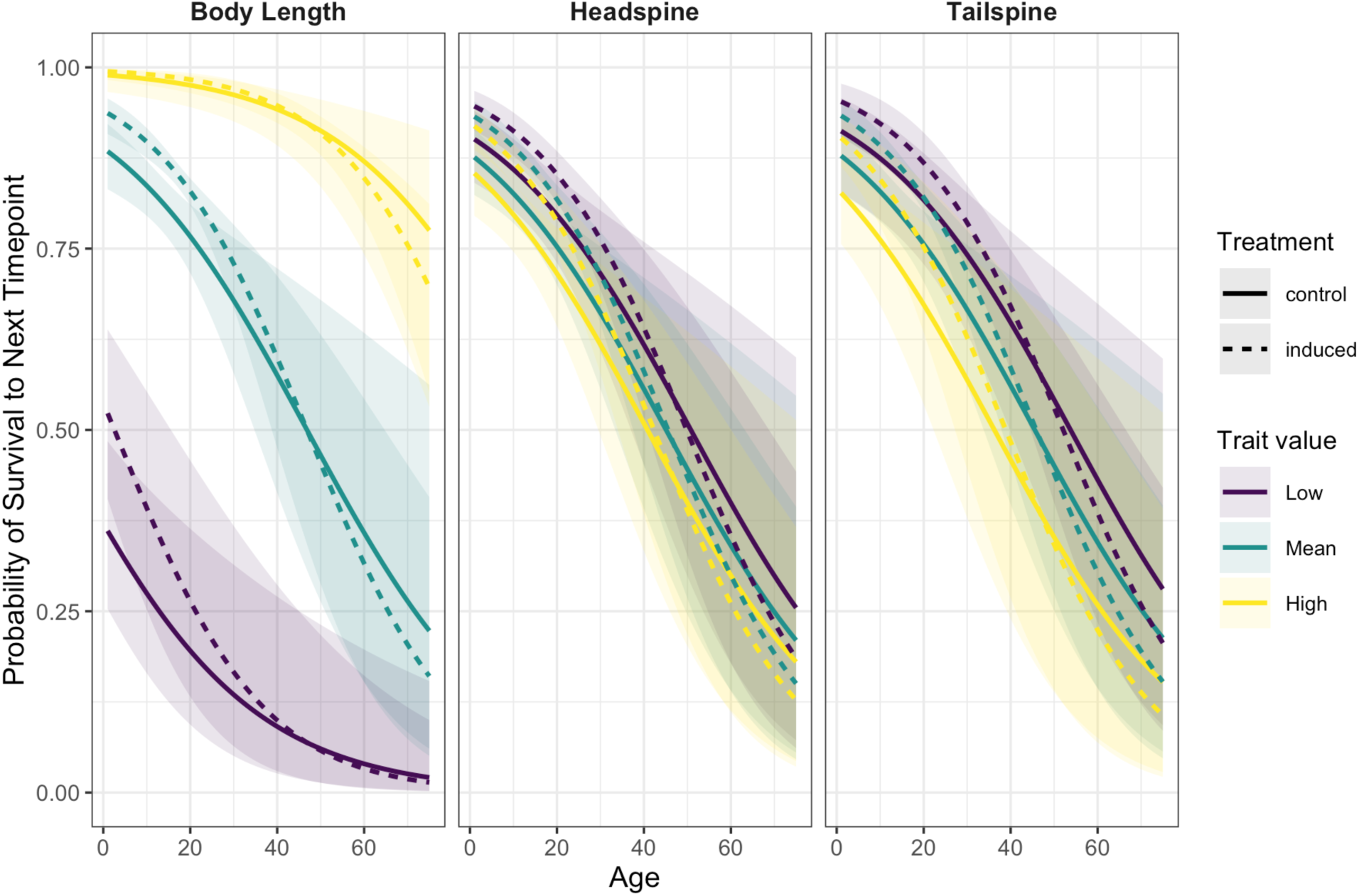
Survivorship and morphological trait distributions do not support size-selective mortality as a driver of morphological differences. Modeled probability of survival to the next timepoint as a function of age, body length, spine length, and treatment, based on a generalized linear model. Survival probability was predicted at 10th (low - purple), 50^th^ (mean – blue), and 90^th^ (high-yellow) percentiles of trait values, with dashed lines representing induced/treated animals and solid lines representing control. Body length values are associated with a greater likelihood of survival at most ages, whereas different spine lengths show minimal effect on predicted survival, regardless of treatment. This suggests that while body length does relate to survival probability in the model, size-selective mortality is not strong enough to cause large differences in defensive morpholgies.

#### Sample Size Considerations

Due to attrition and differences in fecundity and survivorship across 6 generations, the F5 generation suffered from lower sample sizes comparatively, particularly in the control animals. Although we employ Bayesian hierarchical models, which are better equipped to handle unbalanced designs and sparse sample sizes, we conducted a retrospective power analysis using the mean effect size from the F3 and F4 generations (Supplemental Table 6) using the `pwr` R package and the `pwr.t.test` function (Champely et al., 2020).

### Statistical Software and Reproducibility

All statistical analyses and visualizations were conducted in **`R`** (R Core Team, 2021). Data manipulation and cleaning were performed using the **`dplyr`** package (Wickham et al., 2022), and visualizations were generated using `ggplot2` (Wickham, 2016), with additional formatting and layout support from `patchwork` (Pedersen, 2019), `gghighcontrast` (Grosenbach, 2021), and `Rmisc` (Hope, 2022) for summary statistics. Estimated marginal means and post hoc comparisons were calculated using `emmeans` (Lenth & Piaskowski, 2017).

All analyses were conducted using `Rstudio` (Posit Team, 2025) in `RMarkdown` (Allaire et al., 2019) and `Quarto` (Allaire et al., 2025), ensuring full reproducibility of code, figures, and statistical outputs. Annotated Quarto files and datasets are available on GitHub at https://github.com/shannonsnyder/daphnia-transgenerational-plasticity.

## Results

### Defensive and Somatic Traits Display Distinct Ontogenetic Patterns

*Daphnia lumholtzi* exhibited distinct developmental trajectories across their lifespan for body length, headspine, and tailspine structures, regardless of treatment (Figure 3). Defensive spine traits displayed age-dependent plasticity, with early-life elongation followed by gradual reduction in spine length across all individuals, independent of treatment (Fig. 3B-C). Headspine length peaked (∼850 μm) within the first 10 days and declined steadily to ∼400 μm in older individuals (Figure 3B). Tailspine length reached a delayed maximum (∼1,800 μm) around 20 days before declining to values comparable to those observed in early developmental stages (∼700 µm) (Figure 3C). These general developmental patterns provide the ontogenetic context within which treatment-specific differences (Fig. 4) should be interpreted.

### Predator cue exposure results in an increase in headspine and tailspine length in the exposed generation

Because most morphological changes occur within the first 20 days of life, before spine growth plateaus (Figure 3), we modeled treatment effects on individuals less than 20 days of age. Consistent with previously documented within-generation plastic responses in other species, F0 *Daphnia lumholtzi* reared in predator cue environments exhibited robust and statistically significant increases in both headspine and tailspine length relative to controls, indicating postembryonic predator cue exposure was sufficient to produce a substantial increase in both headspine and tailspine length. In the F0 generation, with predation cues provided from the first post-embryonic day, Bayesian models estimate significant increases in induced defense trait lengths (Figure 4; Table 2). Specifically, we report an estimated increase in headspine length of nearly 70 μm (95% CI: [1.67,132.90]). These differences correspond to a roughly 10% increase in headspine length in exposed Daphnia over control Daphnia. Tailspine length also increased dramatically in the exposed F0 individuals, by nearly 100 μm in directly exposed Daphnia (95% CI: [15.35,181.54]), also representing a roughly 10% increase.

### Defensive Traits Are Inherited Through F4

Phenotypic differences persisted for multiple generations after removal from the inducing environment, in generations sufficiently removed from embryonic and germline exposure. When F0 mothers are exposed, their developing F1 embryos are directly exposed in the brood chamber, and the germline cells within F1 embryos that will produce F2 are also exposed. Thus, F2 represents the last generation with potential germline exposure. Only by the F3 generation are any observed effects transmitted through purely non-genetic mechanisms, as F3 individuals and their germ cells had no direct contact with predator cues.

Despite two intervening generations without exposure, F3 individuals with ancestral predation exposure in F0 exhibited a significant increase in headspine length. Bayesian inference yielded an effect size of almost 40 μm (95% CI: [3.10,74.71]). This represents ∼56% retention of the F0 headspine response magnitude (38.19 μm vs. 68.54 μm). In contrast, tailspine length in F3 showed no significant difference between treated and control individuals (β = 5.81 μm, 95% CI: [-30.24, 41.81]), indicating trait-specific attenuation of the tailspine below a statistically detectable threshold.

In the F4 generation, also completely unexposed to predator cues, both headspine and tailspine lengths were again significantly elevated in the induced animals as compared to the control individuals. Ancestral predator exposure increased the headspine length by 44.36 μm (95% CI: [3.99,84.58]) and the tailspine length by 85.57 μm (95% CI: [16.95,154.28]). Therefore, F4 headspine retained ∼65% of the F0 response magnitude (44.36 vs. 68.54 μm), while F4 tailspine retained ∼86% (85.57 vs. 99.71 μm), demonstrating trait-specific patterns of transgenerational persistence (Table 2). Tailspine showed a distinct pattern from headspine: non-significant in F3 but strongly significant in F4, suggesting potential generational fluctuations in trait expression or differential transgenerational mechanisms between the two defensive structures.

By the F5 generation all measurable differences between induced and control individuals had disappeared. We report an estimate of −7.07 (indicating treated animals have a shorter headspine; 95% CI: [-51.23,36.78]) for the headspine and −19.21 μm (95% CI: [-77.17, 37.50] for tailspine length (Table 2, Figure 4). Estimated differences were small with 95% credible intervals widely overlapping zero for both headspine (95% CrI: [-51.23, 36.78]) and tailspine (95% CrI: [-77.17, 37.50]), indicating no detectable treatment effect. These findings indicate that the induced phenotypes ultimately reverted to a statistically undetectable baseline in the absence of continued environmental cues over 5 unexposed generations. Attrition across six generations resulted in a small sample size for this generation, making it difficult to discern if we lack the statistical power to detect a subtle effect.

Model fit was strong across all analyses, with marginal R² values ranging from 56-66% for headspine models and 94-98% for tailspine models, indicating that treatment and body length together explain substantial variance in spine morphology. Frequentist linear mixed-effects models yielded highly consistent results (Supplemental Table 2), with overlapping confidence intervals and similar effect size estimates, confirming the robustness of our inferences across statistical frameworks.

These results provide rare empirical evidence of vertebrate predator-induced transgenerational plasticity persisting through the F4 generation. The temporal trajectory of treatment effects across generations (Figure 4) reveals both the persistence and ultimate decay of transgenerational plasticity. Headspine effects declined from F0 through F5, while tailspine showed an unexpected resurgence in F4 after non-significant F3 effects, highlighting trait-specific dynamics. The persistence of these effects through four generations, particularly in a species with rapid generation times (∼2 weeks), demonstrates substantial temporal stability of non-genetic inheritance.

### Treatment Effects on Body Length are Minimal

In contrast to defensive traits, body length increased rapidly as the animals aged in both control and induced individuals, reaching an early asymptote (∼2,500 μm) around 25 days. This growth pattern was strongly correlated with age (Pearson’s r = 0.85; Supplemental Figure S1). Visual inspection of ontogenetic patterns (Figure 3) showed overlap between body length trajectories between control and induced animals, likely demonstrating trait-specific inheritance of defensive morphologies independent of general body size effects. Indeed, confidence intervals for the Bayesian models illustrating the effect of treatment on body length across generations largely overlap zero, with probability of direction (PD) estimates hovering around 50% (PD > 97.5% typically indicates strong directional evidence), and estimated body size differences between −2.63 and 0.46 μm (treated - control; Table 3), except for the F4 generation. Notably, F4 confidence intervals narrowly overlapped zero (−122.54, 2.22) with an estimated body length difference of −61.45 μm, and a PD of 97.1%, just shy of the conventional 97.5%, providing some support for a smaller body size due to predation exposure in this generation only. Effect sizes were negligible across most generations (Cohen’s d = −0.05 to 0.03), with only F4 showing a small to medium effect (d = −1.16), further supporting that ancestral predation exposure primarily affects defensive structures rather than general body size.

Together, these ontogenetic analyses demonstrate that transgenerational inheritance of predator-induced traits is largely specific to defensive structures and occurs independently of changes in body size. There is, however, some evidence of smaller body sizes due to ancestral predation exposure in the F4 generation alone.

These results demonstrate that predator-induced morphological plasticity can be inherited through at least four generations and is largely concentrated to defensive spines rather than body length. This inheritance is finite and trait-dependent, with detectable headspine elongation through the F4 generations. The decrease of effect sizes and the disappearance of measurable trait differences in F5 underscores the transient nature of this plasticity and suggests that the mechanism of transgenerational epigenetic inheritance decays over time without environmental cue reinforcement (Figure 4).

#### Differences in Growth Rate and Trait Distributions Do Not Explain Induced Trait Expression

To evaluate whether the observed morphological differences between induced and control individuals could be attributed to variation in growth dynamics (rather than final trait size) or trait distributions, we assessed whether treatment effects could be explained by differences in developmental timing, size-selective mortality, or trait-based survival.

The most pronounced morphological changes occurred early in life, prior to 20 days of age (Figure 3). This period includes the onset of reproduction (typically 7–12 days), during which *D. lumholtzi* mothers are not yet fully grown. Morphological change peaked around this time for all traits, particularly for body length and total body size (Figure 5).

Importantly, no significant differences in growth rate were detected between treatment groups for any of the four traits examined (body length, headspine, tailspine, and total body size), indicating that predator exposure did not alter the rate of somatic or defensive trait development (Table 4). Specifically, treatment effects were non-significant for body length (Est. = 5.70 ± 7.94 μm, F = 0.52, p = 0.47), total body size (Est. = 9.87 ± 19.20 μm, F = 0.27, p = 0.61), headspine (Est. = 6.51 ± 4.57 μm, F = 2.06, p = 0.15), and tailspine (Est. = 1.88 ± 7.57 μm, F = 0.06, p = 0.80). Generation-specific analyses similarly revealed no treatment effects on growth rates in any individual generation (all p > 0.05; Supplemental Table 4), confirming that differences in defensive trait expression persist consistently across F0-F5 independent of developmental timing. These results suggest that the observed morphological differences are not due to accelerated or delayed growth in induced individuals but instead reflect differences in final trait expression independent of growth rate.

**Table 4:**
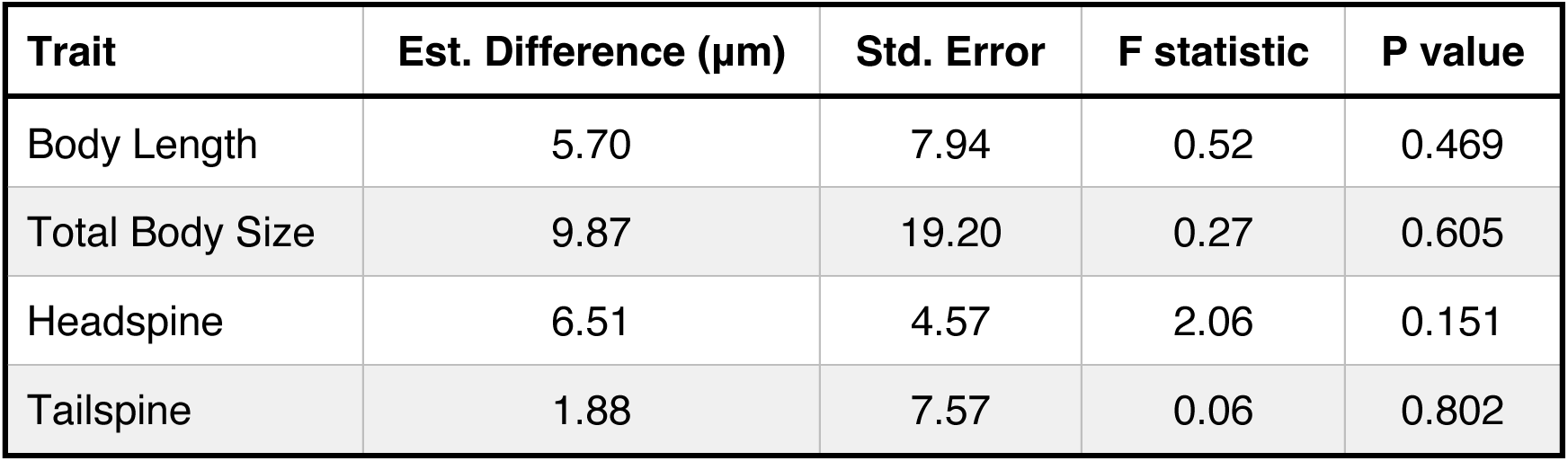
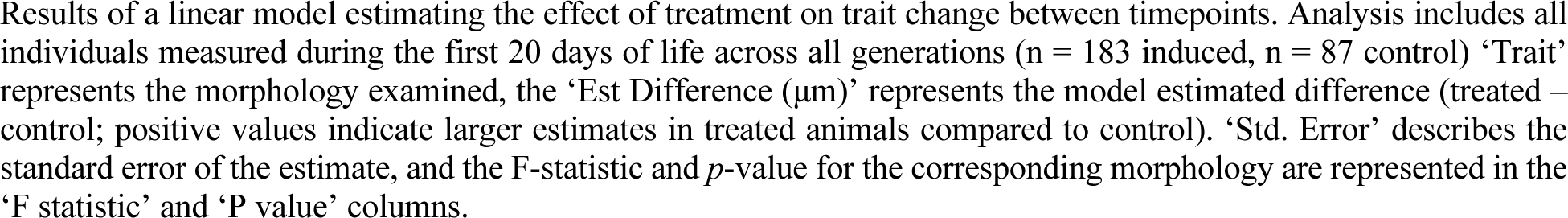
Growth rate is not influenced by treatment.

| Trait | Est. Difference ($\mu\text{m}$) | Std. Error | F statistic | P value |
| --- | --- | --- | --- | --- |
| Body Length | 5.70 | 7.94 | 0.52 | 0.469 |
| Total Body Size | 9.87 | 19.20 | 0.27 | 0.605 |
| Headspine | 6.51 | 4.57 | 2.06 | 0.151 |
| Tailspine | 1.88 | 7.57 | 0.06 | 0.802 |

Results of a linear model estimating the effect of treatment on trait change between timepoints. Analysis includes all individuals measured during the first 20 days of life across all generations ( $n = 183$ induced, $n = 87$ control) 'Trait' represents the morphology examined, the 'Est Difference ( $\mu\text{m}$ )' represents the model estimated difference (treated – control; positive values indicate larger estimates in treated animals compared to control). 'Std. Error' describes the standard error of the estimate, and the F-statistic and $p$ -value for the corresponding morphology are represented in the 'F statistic' and 'P value' columns.

To assess whether trait distributions might reflect selective mortality, such as the preferential survival of individuals with longer spines, we compared growth trajectories of survivors and non-survivors during the first 20 days of life. Plotting developmental trajectories for body length, headspine residuals, and tailspine residuals (residuals control for body size) by survival status (Figure 6) showed that while survivors tended to have a larger body length, **residual spine lengths showed overlapping distributions and trajectories** between survivors and non-survivors. Overlapping distributions of spine residuals between survivors and non-survivors (Figure 6) indicate that trait-based selection is unlikely to account for the observed morphological differences.

The absence of an effect of spine length on survivorship was further supported by a generalized linear model (GLM) predicting survival status from body length, headspine, tailspine, age, and treatment. The model revealed that **body length was positively associated with survival** (β = 0.0030, *p* < 0.001), while **headspine and tailspine lengths were not significant predictors** (headspine: β = –0.0010, *p* = 0.058; tailspine: β = –0.0007, *p* = 0.110). Individuals from predator-exposed lineages showed significantly higher survival than controls (β = 0.678, p < 0.001), suggesting that transgenerational defensive trait expression may confer fitness benefits even in the absence of direct predator exposure. Importantly, this survival advantage was independent of spine length, as neither headspine nor tailspine significantly predicted survival status. The model explained a substantial portion of deviance (ΔDeviance = 328.8; AIC = 2554.1), but **no evidence emerged for size-or trait-selective mortality based on spine length** (Figure 7).

### Power Analysis

F5 sample sizes were reduced due to natural attrition across six generations (n=8 control individuals, 41 induced; Table 1). Retrospective power analysis using mean effect sizes from F3-F4 generations yielded estimated power of 0.30 for headspine and 0.66 for tailspine (Supplemental Table 5). These estimates are conservative because they do not account for repeated measures (average 12.2 observations per individual for induced, 5.6 for control; Table 1), which increase effective sample size. While we cannot definitively distinguish between trait decay and insufficient power in F5, the pattern across F0-F5 suggests gradual attenuation consistent with temporal decay of transgenerational effects.

These findings demonstrate that transgenerational plasticity in *D. lumholtzi* operates primarily on defensive structures, persists for at least four generations following ancestral exposure, and is not due to size-selective or trait-selective mortality.

## Discussion

Our findings demonstrate that predator-induced morphological defenses in *Daphnia lumholtzi* persist through at least four unexposed generations via non-genetic mechanisms. Critically, measurable effects in F3, F4, and (to a lesser extent) F5 occurred in cohorts with no direct, embryonic, or germline exposure to predator cues, a necessary criterion for distinguishing transgenerational plasticity from extended maternal effects (Agrelius & Dudycha, 2025; Skinner, 2008). This temporal persistence is notable given that F3 represents the first generation where developing germ cells had no contact with the inducing signal, and F4-F5 are even further removed. These data contribute to a limited but growing body of evidence demonstrating that environmentally induced phenotypes can be inherited across multiple generations through epigenetic or other non-genetic mechanisms.

Our experimental design intentionally employed a single clonal lineage to isolate non-genetic effects from standing genetic variation, enabling us to definitively demonstrate transgenerational inheritance independent of allelic differences. However, this design limits inference to one genetic background. The existence of ‘uninducible’ *D. lumholtzi* clones with constitutive defenses (Engel & Tollrian, 2009), even from nearby populations in Arizona, demonstrates substantial genetic variation in plasticity and defense strategies within this species. Future work must examine whether the 4-generation persistence we observed is representative across clones or population-specific, and whether genotype-by-environment interactions influence the temporal dynamics of transgenerational effects (Auld et al., 2010; Scoville & Pfrender, 2010).

### Induced Phenotypes Persist Across Generations but Fade Without Cues

Headspine and tailspine elongation persisted through four unexposed genetically identical generations, with measurable effects in F3 and F4 despite the absence of continued induction. This transgenerational persistence supports the hypothesis that environmental information can be retained and expressed across generations via non-genetic mechanisms (Gapp et al., 2014; Heard & Martienssen, 2014; Rechavi et al., 2014).

By the F5 generation, trait differences between induced and control individuals were not detectable. This regression to baseline aligns with evolutionary expectations that plastic traits fade when environmental cues vanish, minimizing fitness costs associated with unnecessary defenses (DeWitt et al., 1998; Van Buskirk & Steiner, 2009).

Headspine and tailspine displayed divergent transgenerational trajectories, suggesting trait-specific non-genetic regulation. Headspine effects declined rather monotonically across generations (68 μm → 38 μm → 44 μm → −7 μm), consistent with gradual erosion of epigenetic marks. In contrast, tailspine showed non-significant effects in F3 (6 μm) but pronounced effects in F4 (86 μm), approaching F0 magnitude (100 μm). This “resurgence” pattern could reflect: (1) differential stability of epigenetic marks regulating the two traits, (2) compensatory developmental mechanisms favoring tailspine expression when headspine inheritance weakens, or (3) stochastic fluctuations in small sample sizes. These trait-specific dynamics highlight that transgenerational plasticity is not uniformly expressed across correlated traits and may involve independent regulatory mechanisms (Bonduriansky & Day, 2009; Herman & Sultan, 2011; Uller et al., 2013). Future work should examine whether these two defensive structures are regulated by distinct epigenetic pathways.

Retrospective power analysis for F5 yielded estimated power of 0.30 for headspine and 0.66 for tailspine, based on mean effect sizes from F3-F4 generations. While we cannot definitively distinguish between true decay and insufficient power in F5, several lines of evidence support the former: (1) the decline in effect sizes across F0-F4, (2) credible intervals for F5 estimates centered near zero rather than showing directional trends, and (3) the biological expectation that transgenerational effects should eventually attenuate without environmental reinforcement. Achieving high statistical power in ecological studies, especially with multigenerational designs is inherently challenging (Kimmel et al., 2023) but our use of repeated measures and hierarchical modeling maximized inferential strength given practical constraints.

The 5-generation persistence of predator-induced defenses has implications for population dynamics in variable predation regimes. In systems with fluctuating predator densities (seasonal fish migrations, predator reintroductions, reproductive timing), prey populations may remain partially defended for 4-5 generations (several months in this *Daphnia* species) after predator removal. This “ecological memory” could buffer populations during temporary predator absences but may incur costs if defenses persist longer than threats. Fitness consequences, including fecundity and survival across generations will be the focus of future work in our predator-prey system.

### *Daphnia lumholtzi* Induced Defenses Are Robust, Age-Specific, and Not Driven by Selection

Exposure to predator cues in the F0 generation triggered strong morphological defenses in *Daphnia lumholtzi*, with significant elongation of both headspine and tailspine structures. Post-embryonic exposure alone was sufficient to induce these traits in our Arizona clone, demonstrating that embryonic priming is not universally required for defense induction in this genus (Laforsch & Tollrian, 2004). This finding underscores the differences in inducibility between clones and species and highlights the need to assess transgenerational inheritance in a variety of systems (Auld et al., 2010; Scoville & Pfrender, 2010; Shahmohamadloo et al., 2025). This is further illustrated by the characterization of “uninducible” *D. lumholtzi* clones which constitutively express defense phenotypes even in the absence of predation cues (Engel & Tollrian, 2009).

Importantly, these trait differences were not attributable to size-selective or trait-selective mortality. Survivors and non-survivors showed overlapping developmental trajectories, and survival probability was primarily associated with body length, not spine morphology. This confirms that the observed phenotypic differences are induced responses rather than artifacts of differential survivorship among individuals based upon armor morphology (Dieckmann et al., 2006; Kingsolver et al., 2001).

We also found that spine lengths were most prominent in early life and declined with age, a pattern distinct from somatic growth. This suggests stage-specific optimization of defenses, likely targeting periods of highest vulnerability to predation (Nagano & Yoshida, 2020; E. E. Werner & Gilliam, 1984). Smaller, younger Daphnia likely experience higher predation rates from gape-limited fish predators (Nagano & Yoshida, 2020), making early-life defenses particularly adaptive. The age-dependent decline in spine length we observe may represent adaptive reallocation from defense to reproduction as individuals reach body sizes less vulnerable to predation. This ontogenetic trajectory, previously unreported for *D. lumholtzi* headspines, parallels patterns in other inducible defense systems where defenses are most pronounced during vulnerable life stages (Nagano & Yoshida, 2020) and documented tailspine shortening with age in *Daphnia* (Gu et al., 2021).

### Mechanisms of Non-genetic Inheritance and Evolutionary Implications

The persistence and gradual decline of induced traits across generations strongly suggests epigenetic inheritance as a likely mechanism. Having established the phenotypic baseline and generational persistence, future mechanistic work can now uncover the molecular mechanisms underlying these patterns. Candidate mechanisms include, DNA methylation, histone modification, and chromatin-level changes that facilitate the transmission of plasticity without altering the genetic code (Banerjee & Roychoudhury, 2017; Casadesús, 2016; Davidowitz, 2009; Ho & Burggren, 2010; Li & Zhang, 2014; Niederhuth et al., 2016; Robichaud et al., 2012; M. S. Werner et al., 2023; Yaniv, 2014). The temporal decay pattern we observed is consistent with incomplete erasure of epigenetic marks (Dias & Ressler, 2014; Gapp et al., 2014; Grishkevich & Yanai, 2013), as documented in other invertebrate and vertebrate systems including *Caenorhabditis elegans* (Kaneshiro et al., 2019), the asexual freshwater snail, *Physa acuta* (Dejeux et al., 2025), *Daphnia magna* (Asselman et al., 2016; Vandegehuchte, De Coninck, et al., 2010), as well as vertebrate systems including mice (Carone et al., 2010) and humans (Mulligan et al., 2025). By identifying these generational phenotypic dynamics, we establish a foundation for uncovering the molecular mechanisms underlying generation-specific adaptive phenotypes, a key priority for future research.

Our focus on vertebrate predation addresses an important gap in transgenerational plasticity research. While numerous studies examine *Daphnia* responses to invertebrate predators such as *Chaoborus* larvae (Agrawal et al., 1999; Horstmann et al., 2022), studies of vertebrate predator-induced transgenerational effects remain scarce (Tariel et al., 2020). This gap is consequential because vertebrate predators typically exert more consistent selective pressure than invertebrate predators (Batabyal, 2023; Wooster, 1994) and are central to current conservation efforts. Our findings have direct relevance for predator reintroduction and rewilding efforts, where prey populations may exhibit defenses for 4-5 generations (∼2-3 months in *D. lumholtzi*) following ancestral exposure. Understanding the temporal dynamics of these transgenerational responses, especially in a keystone aquatic species, could improve predictions of predator-prey dynamics and ecosystem restoration outcomes (Bracis & Wirsing, 2021; Delibes-Mateos et al., 2019).

Our findings align with theoretical predictions that transgenerational plasticity should evolve when parental and offspring environments are positively correlated (Agrawal et al., 1999; Herman & Sultan, 2011). The 4-generation persistence we observe (∼2-3 months in this system) matches timescales over which predator densities fluctuate in natural lakes due to seasonal migrations, overwinter mortality, or breeding events. However, the eventual decay by F5 is also adaptive as prolonged expression of costly defenses in predator-free environments would reduce fitness through allocation trade-offs (DeWitt et al., 1998; Van Buskirk & Steiner, 2009). The temporal dynamics we document therefore suggest fine-tuning of inheritance duration to match ecological predictability, though testing this hypothesis would require comparative studies across populations experiencing different predation regimes.

Our findings underscore the evolutionary relevance of environmental history in the absence of genetic variation (Jablonka & Lamb, 2006; Laland et al., 2015; Pigliucci & Müller, 2010). Transgenerational plasticity allows organisms to respond to ancestral conditions, potentially buffering populations against short-term environmental fluctuations (Donelson et al., 2018; Uller et al., 2018). Because our study intentionally used a single *Daphnia* population, future work should explore responses across diverse genotypes (Auld et al., 2010). The existence of “uninducible” clones with constitutive defenses raises intriguing questions about genetic assimilation and the long-term fate of plastic traits (Lande, 2009; West-Eberhard, 2003).

### Conclusions

This work advances understanding of transgenerational plasticity in a keystone aquatic genus by documenting vertebrate predator-induced morphological effects in key generations sufficiently removed from embryonic and germline exposure, therefore revealing trait-specific temporal persistence in defensive structures. Our findings align with previous work documenting clutch size persistence in *D. ambigua* (Walsh et al., 2014), helmet length inheritance in *D. cucullata* (Agrawal et al., 1999), and multi-generational erosion of plastic traits in *D. pulicaria* (Landy et al., 2024), while crucially extending the temporal window of observation for predator-induced effects to key, previously undocumented generations.

This transgenerational plasticity may buffer populations against short-term ecological fluctuations, while its eventual attenuation minimizes the costs of unnecessary trait expression (Herman & Sultan, 2011; Uller et al., 2013). Future research should integrate molecular profiling (DNA methylation, chromatin modifications, differential transcription) across F0-F5 generations with multi-clone comparative studies to determine both the mechanistic basis and generality of these transgenerational effects. Such integrative approaches will advance understanding of how organisms respond to environmental variation across generational timescales and the role of non-genetic inheritance in evolutionary adaptation to changing environments.

## Data availability statement

The raw data and full analysis scripts that support the findings of this study are openly available in Markdown on GitHub (https://github.com/shannonsnyder/daphnia-transgenerational-plasticity).

## Author contributions

**Conceptualization and Methodology:** SNS and WAC conceived and designed the study. **Investigation:** SNS, WCM, and SLJ reared the animals and collected the imaging data. **Data Collection and Curation:** SNS, EBC and TB curated the morphological measurements. **Formal Analysis:** SNS and WAC analyzed the data. **Visualization:** SNS created the data visualizations. **Writing – Original Draft:** SNS wrote the manuscript. **Writing – Review & Editing:** WAC provided revisions. All authors approved the final manuscript.

## Acknowledgements

We would like to thank M. Lynch for his contribution of the *Daphnia lumholtzi* (SAG) utilized for this study. We would also like to thank Mark Currey for his assistance with stickleback husbandry and the Cresko lab for valuable feedback. Finally, we thank Emily Willams for collection and maintenance of *Daphnia* prior to their arrival in the Cresko lab.

## SUPPLEMENTAL

**Supplemental Figure 1.**
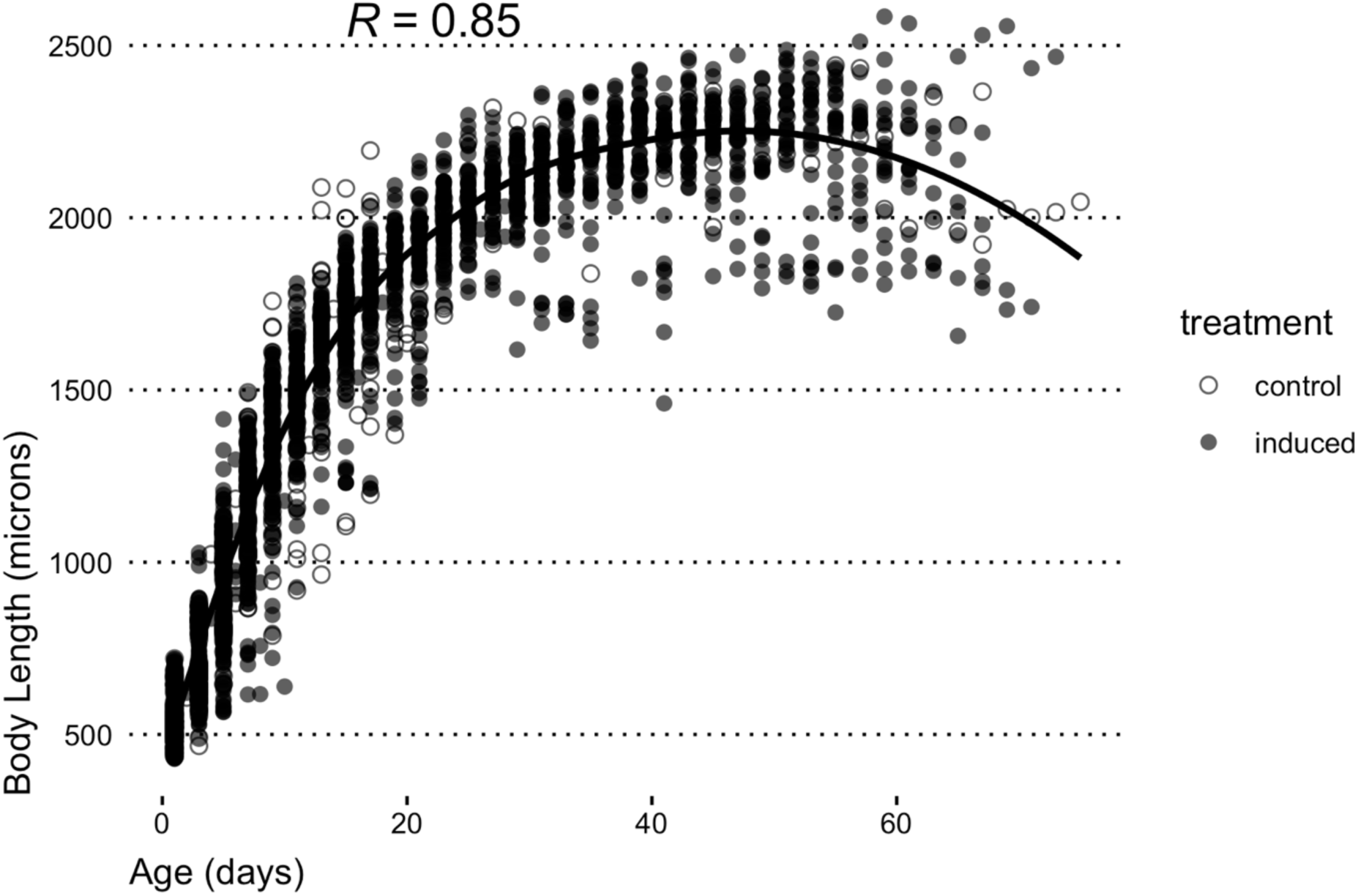
Ontogenetic growth trajectories for Daphnia body length across treatments show high correlation between age and body length. Scatterplot of individual body length measurements (μm) as a function of age (days) for control (open circles) and induced (filled circles) treatment across all generations. The solid line shows the overall fitted growth curve (Pearson’s r =0.85) across all data. Growth patterns are similar between treatment, indicating that environmental induction principally affects defensive traits rather than somatic traits in this clone.

**Supplemental Figure 2:**
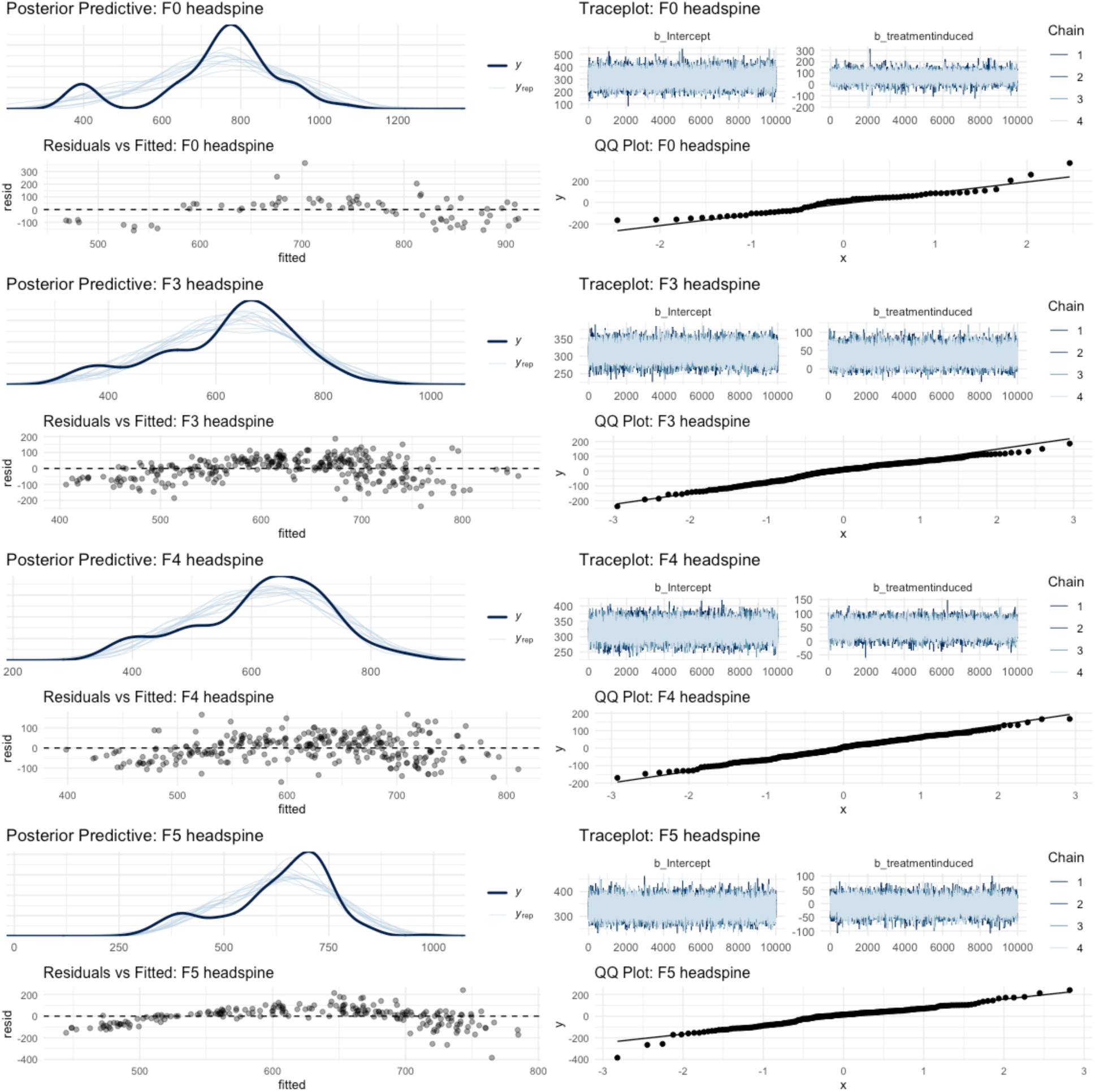
Bayesian model diagnostic plots for the headspine trait across generations. Each row represents an independent model. The panels display posterior predictive checks (model fit), residuals vs. fitted values (homoscedasticity), MCMC traceplots (convergence), and Quantile-Quantile (QQ) plots (normality of residuals). The successful convergence of the MCMC chains and the adequate fit shown by the predictive checks and residual plots support the validity of the models for inference.

**Supplemental Figure 3:**
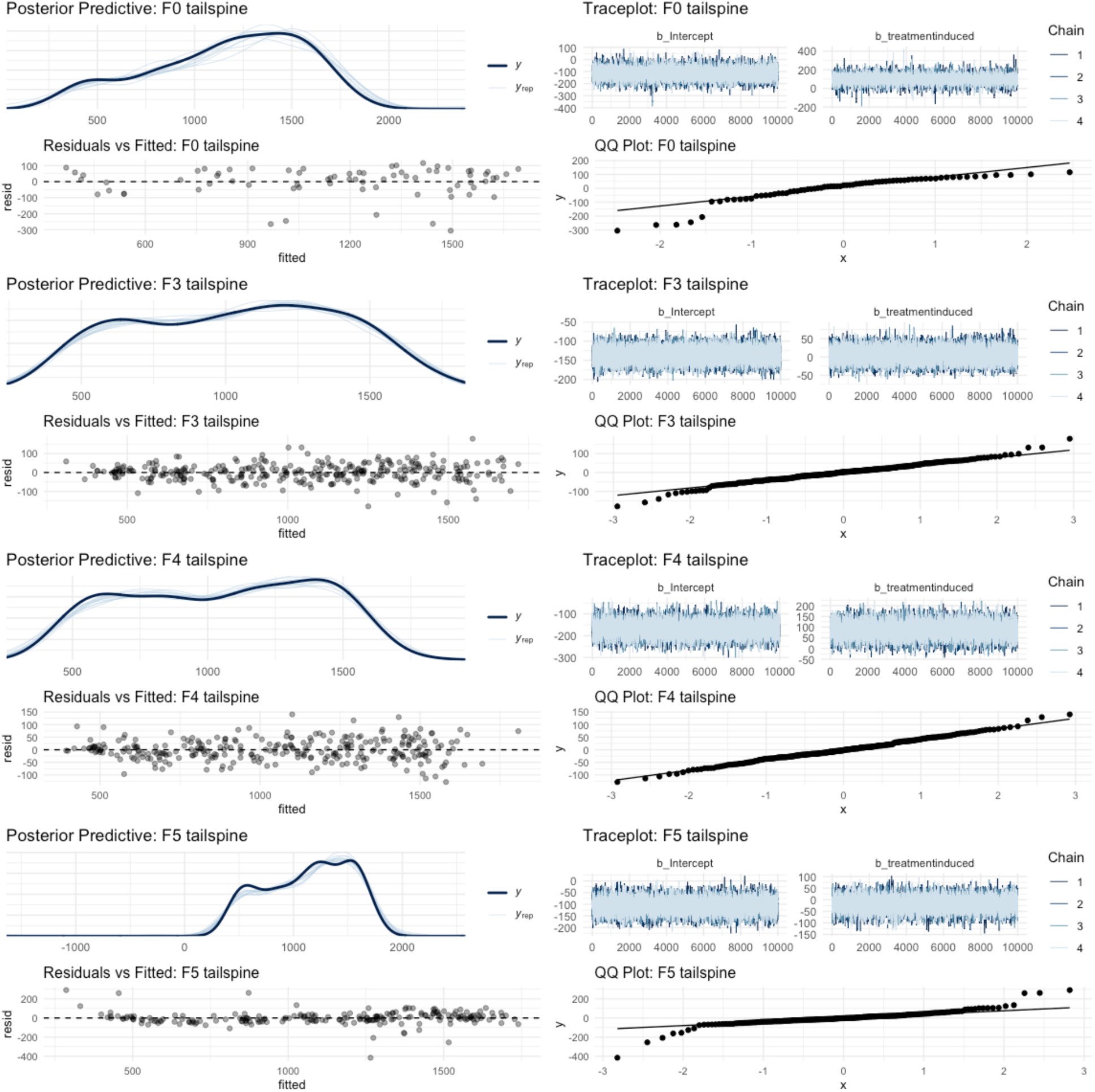
Bayesian model diagnostic plots for the tailspine trait across generations. Each row represents an independent model. Plots include posterior predictive checks (model fit), residuals vs. fitted values (homoscedasticity), MCMC traceplots (convergence), and Quantile-Quantile (QQ) plots (normality of residuals). The posterior predictive checks show good overlap between observed and simulated data, and the traceplots indicate stable, well-mixed chains. The residuals show no major patterns and are approximately normally distributed, supporting the validity of the models.

**Supplemental Figure 4:**
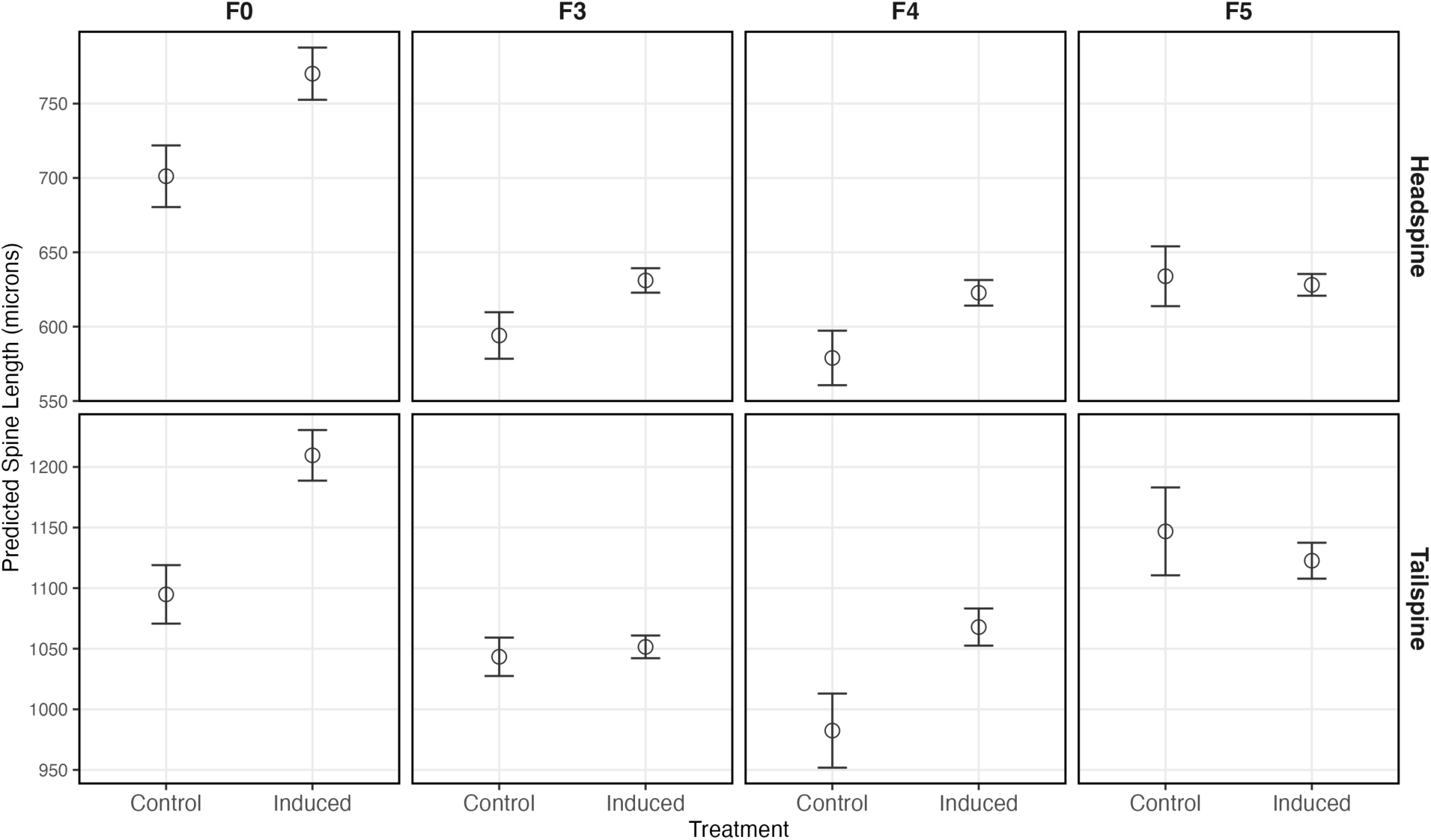
Predicted spine lengths by generation and treatment from linear mixed-effects models. Predicted mean headspine length (top row) and tailspine length (bottom row) for Daphnia in control and induced individuals across generations (F0, F3, F4, F5). Points represent model-predicted means ± standard error from linear mixed-effects models.

**Supplemental Figure 5.**
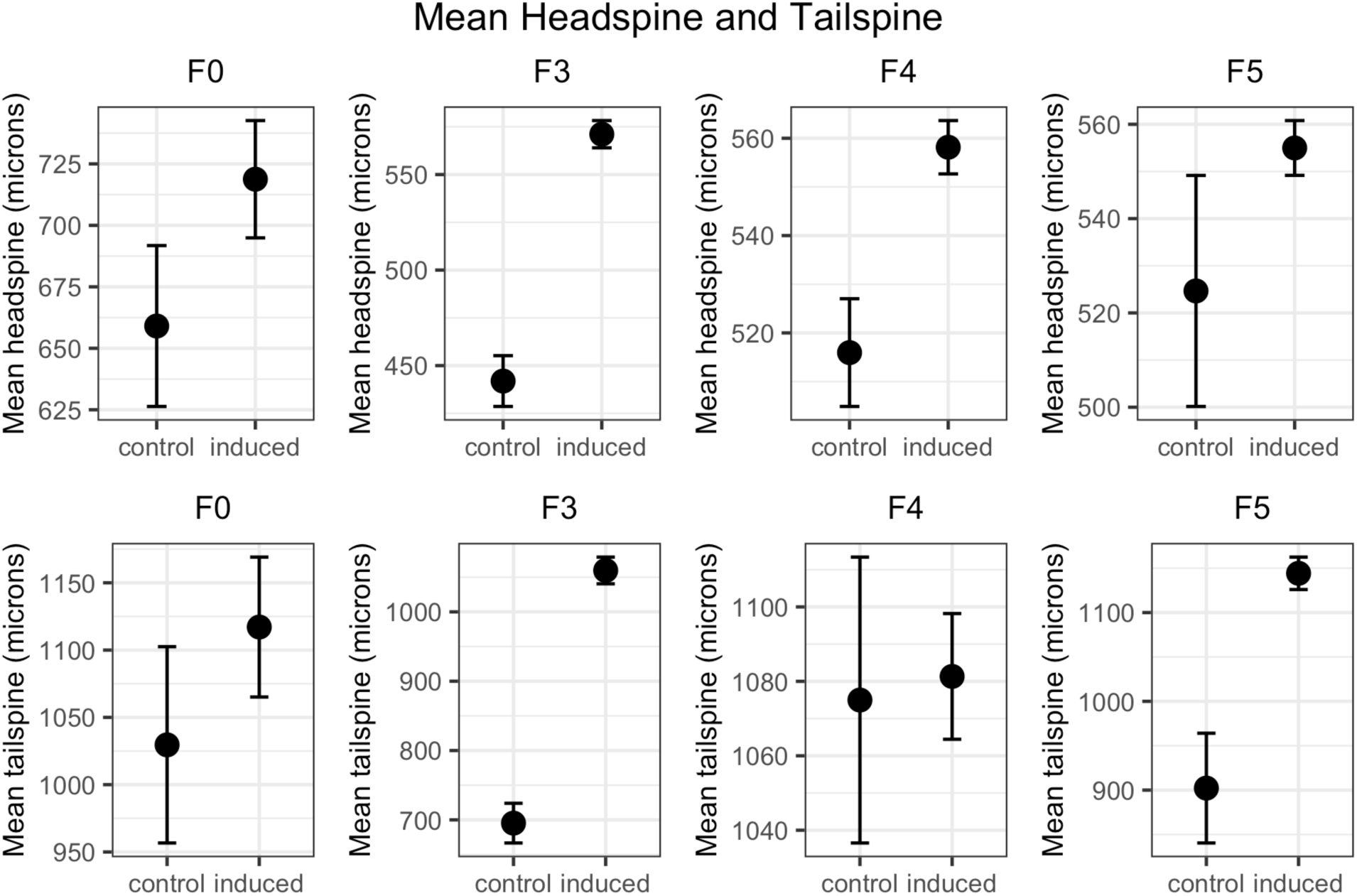
Transgenerational patterns of headspine and tailspine length in Daphnia following ancestral predator cue exposure. Mean (± standard error) headspine (top panels) and tailspine (bottom panels) values are shown for control and induced individuals across four non-exposed generations (F3–F5) and the originally exposed generation (F0). Each panel displays pairwise comparisons between control and induced groups for the indicated generation. Significant induction of both headspine and tailspine length is evident in predator-cue–induced F0 animals, with differential persistence of these defensive traits in subsequent generations. These data illustrate the finite and dynamic pattern of transgenerational plasticity in response to environmental cues. Note: this data is included for transparency, but does not include correction for body sizes, and violates independence assumptions as there are repeated measures.

**Supplemental Figure 6:**
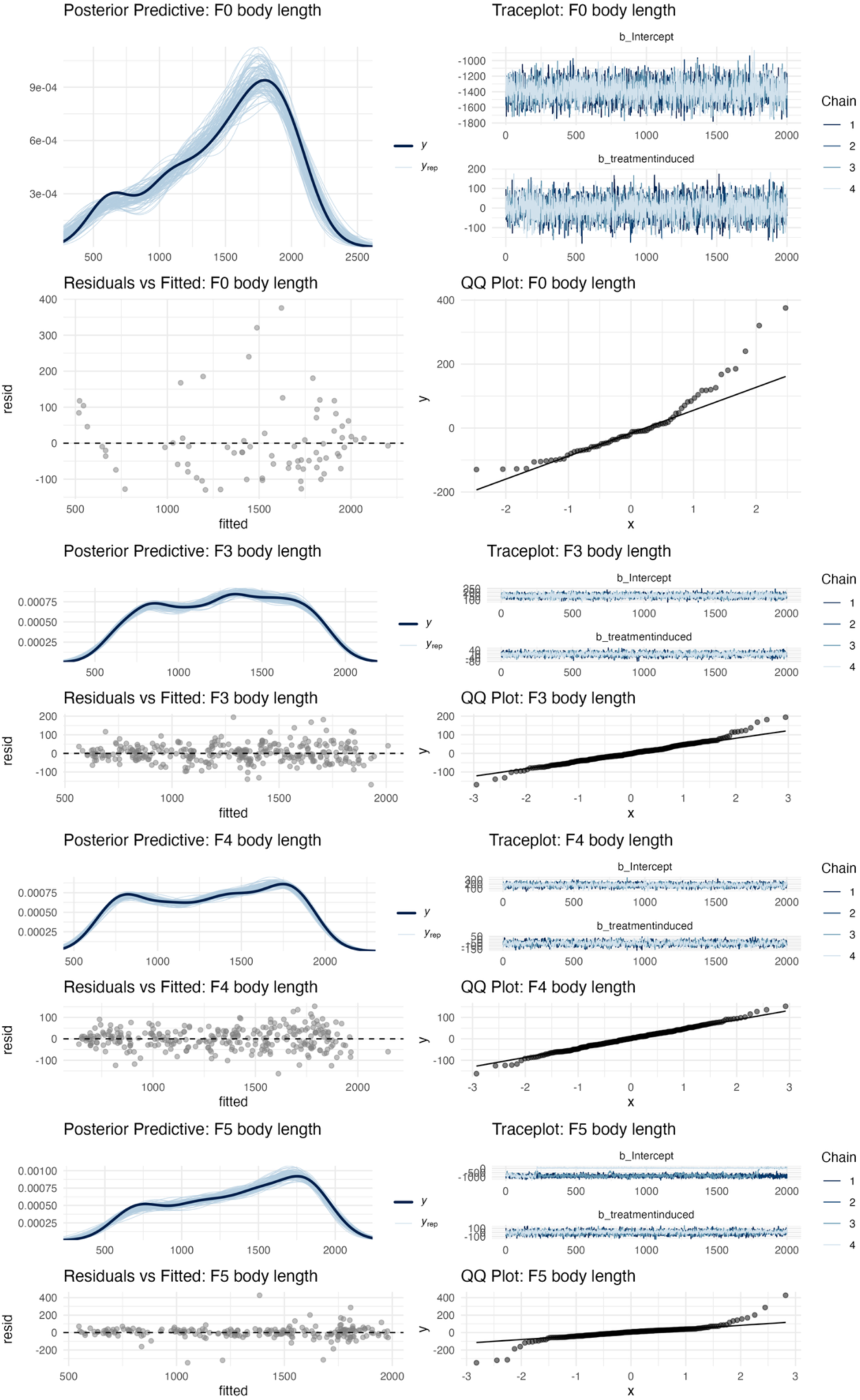
Model diagnostics for body length Bayesian models across generations. For each generation: posterior predictive check (top left), MCMC traceplots (top right), residuals vs fitted (bottom left), and QQ plot (bottom right).

**Supplemental Table 1:**
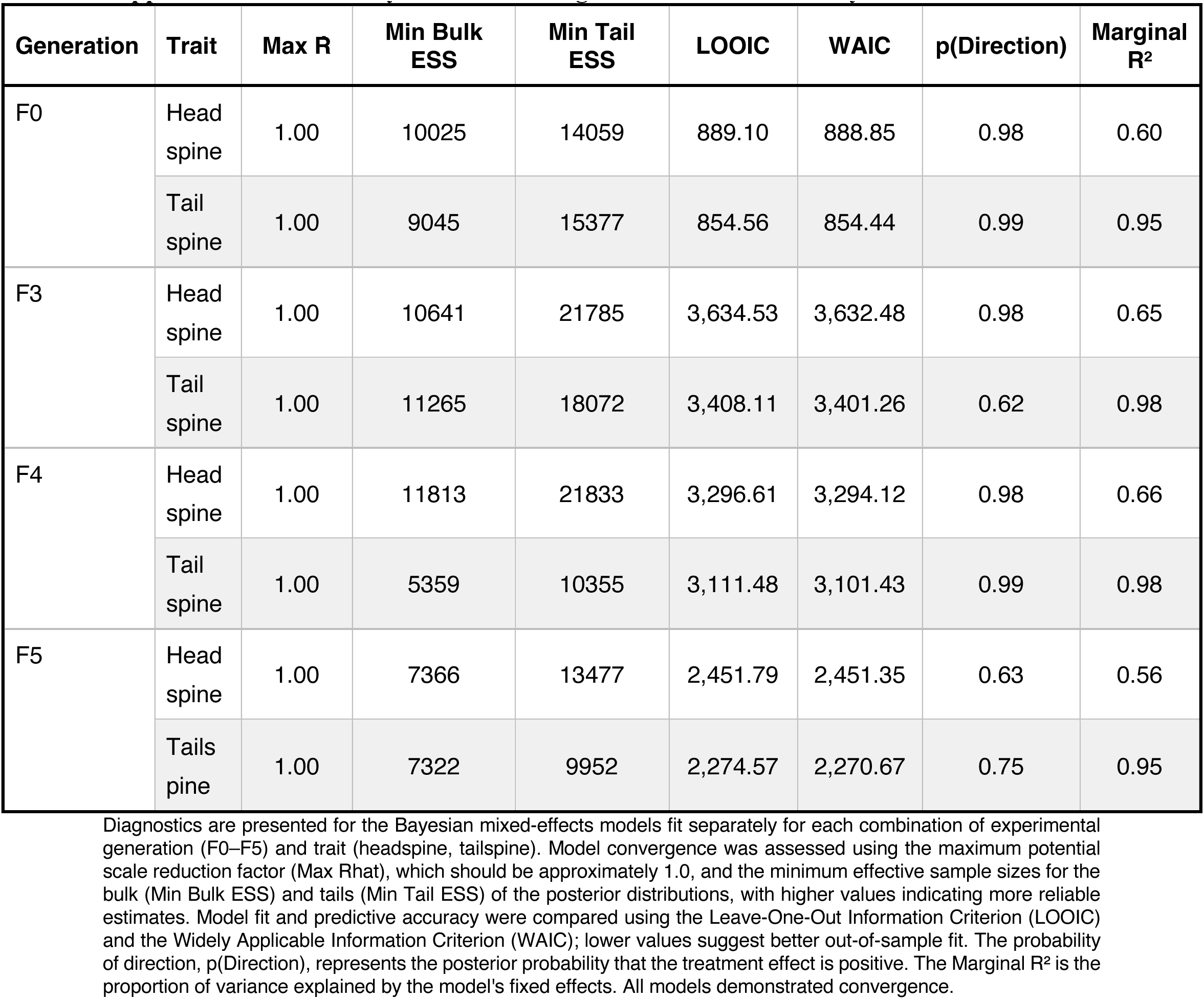
Bayesian Model Diagnostics and Fit Summary.

| Generation | Trait | Max R | Min Bulk ESS | Min Tail ESS | LOOIC | WAIC | p(Direction) | Marginal R<sup>2</sup> |
| --- | --- | --- | --- | --- | --- | --- | --- | --- |
| F0 | Head spine | 1.00 | 10025 | 14059 | 889.10 | 888.85 | 0.98 | 0.60 |
|  | Tail spine | 1.00 | 9045 | 15377 | 854.56 | 854.44 | 0.99 | 0.95 |
| F3 | Head spine | 1.00 | 10641 | 21785 | 3,634.53 | 3,632.48 | 0.98 | 0.65 |
|  | Tail spine | 1.00 | 11265 | 18072 | 3,408.11 | 3,401.26 | 0.62 | 0.98 |
| F4 | Head spine | 1.00 | 11813 | 21833 | 3,296.61 | 3,294.12 | 0.98 | 0.66 |
|  | Tail spine | 1.00 | 5359 | 10355 | 3,111.48 | 3,101.43 | 0.99 | 0.98 |
| F5 | Head spine | 1.00 | 7366 | 13477 | 2,451.79 | 2,451.35 | 0.63 | 0.56 |
|  | Tails pine | 1.00 | 7322 | 9952 | 2,274.57 | 2,270.67 | 0.75 | 0.95 |

Diagnostics are presented for the Bayesian mixed-effects models fit separately for each combination of experimental generation (F0–F5) and trait (headspine, tailspine). Model convergence was assessed using the maximum potential scale reduction factor (Max Rhat), which should be approximately 1.0, and the minimum effective sample sizes for the bulk (Min Bulk ESS) and tails (Min Tail ESS) of the posterior distributions, with higher values indicating more reliable estimates. Model fit and predictive accuracy were compared using the Leave-One-Out Information Criterion (LOOIC) and the Widely Applicable Information Criterion (WAIC); lower values suggest better out-of-sample fit. The probability of direction, p(Direction), represents the posterior probability that the treatment effect is positive. The Marginal R<sup>2</sup> is the proportion of variance explained by the model's fixed effects. All models demonstrated convergence.

**Supplemental Table 2:**
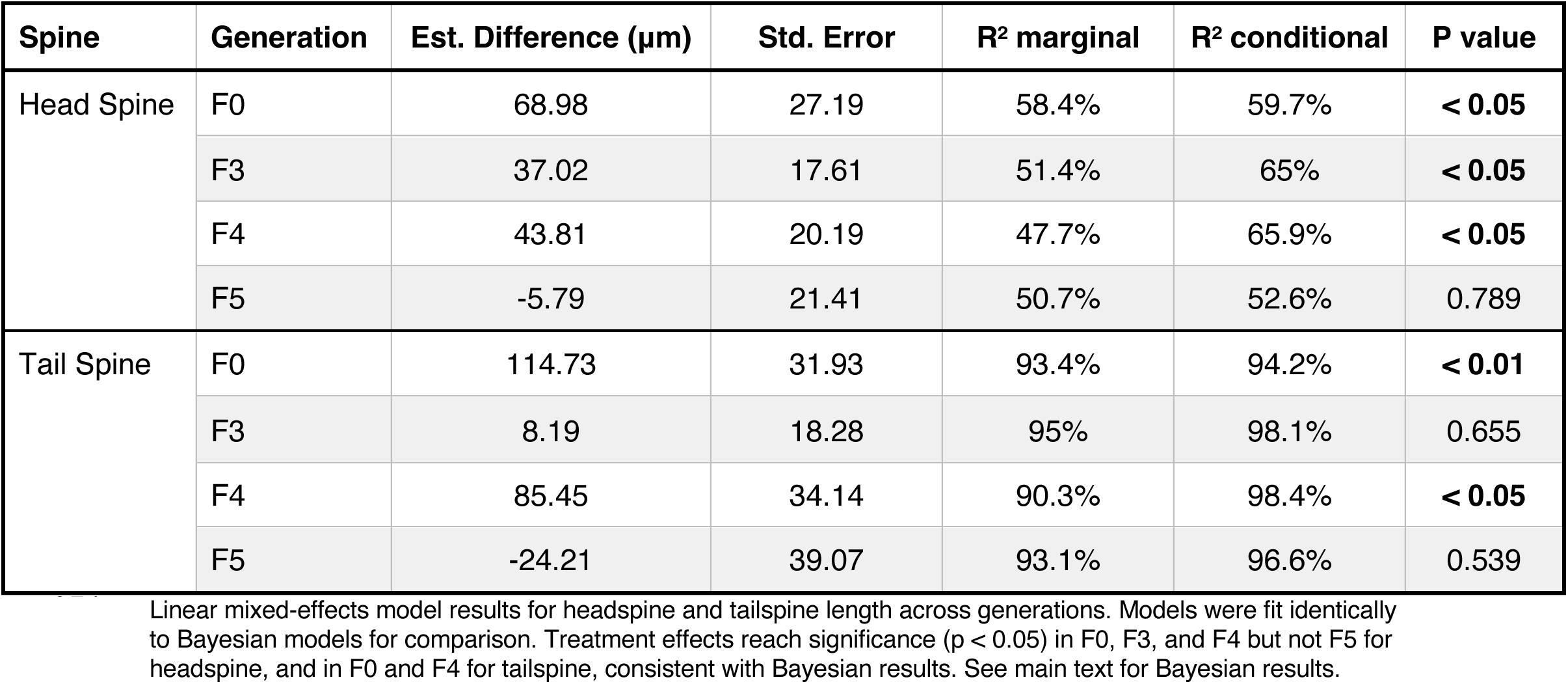
Linear estimates for headspine and tailspine length.

| Spine | Generation | Est. Difference (µm) | Std. Error | R<sup>2</sup> marginal | R<sup>2</sup> conditional | P value |
| --- | --- | --- | --- | --- | --- | --- |
| Head Spine | F0 | 68.98 | 27.19 | 58.4% | 59.7% | < 0.05 |
|  | F3 | 37.02 | 17.61 | 51.4% | 65% | < 0.05 |
|  | F4 | 43.81 | 20.19 | 47.7% | 65.9% | < 0.05 |
|  | F5 | -5.79 | 21.41 | 50.7% | 52.6% | 0.789 |
| Tail Spine | F0 | 114.73 | 31.93 | 93.4% | 94.2% | < 0.01 |
|  | F3 | 8.19 | 18.28 | 95% | 98.1% | 0.655 |
|  | F4 | 85.45 | 34.14 | 90.3% | 98.4% | < 0.05 |
|  | F5 | -24.21 | 39.07 | 93.1% | 96.6% | 0.539 |

Linear mixed-effects model results for headspine and tailspine length across generations. Models were fit identically to Bayesian models for comparison. Treatment effects reach significance ( $p < 0.05$ ) in F0, F3, and F4 but not F5 for headspine, and in F0 and F4 for tailspine, consistent with Bayesian results. See main text for Bayesian results.

**Supplemental Table 3:** Goodness-of-fit and diagnostic statistics for linear mixed-effects models.

| Spine | Generation | Est. (µm) | Std. Error | t-value | Marginal R<sup>2</sup> | Conditional R<sup>2</sup> | AIC | BIC | Log-Likelihood |
| --- | --- | --- | --- | --- | --- | --- | --- | --- | --- |
| Headspine | F0 | 68.98 | 27.19 | 2.54 | 58.4% | 59.7% | 881.0 | 892.5 | -435.5 |
|  | F3 | 37.02 | 17.61 | 2.10 | 51.4% | 65% | 3651.0 | 3669.8 | -1820.5 |
|  | F4 | 43.81 | 20.19 | 2.17 | 47.7% | 65.9% | 3316.0 | 3334.3 | -1653.0 |
|  | F5 | -5.79 | 21.41 | -0.27 | 50.7% | 52.6% | 2454.2 | 2470.9 | -1222.1 |
| Tailspine | F0 | 114.73 | 31.93 | 3.59 | 93.4% | 94.2% | 870.5 | 882.0 | -430.3 |
|  | F3 | 8.19 | 18.28 | 0.45 | 95% | 98.1% | 3485.1 | 3503.9 | -1737.6 |
|  | F4 | 85.45 | 34.14 | 2.50 | 90.3% | 98.4% | 3225.7 | 3244.0 | -1607.9 |
|  | F5 | -24.21 | 39.07 | -0.62 | 93.1% | 96.6% | 2427.1 | 2443.8 | -1208.5 |

Models were fit independently for each combination of experimental generation (F0, F3, F4, F5) and morphological trait (headspine, tailspine). The estimate represents the effect of treatment in µm. Std. error indicates the standard error, and the t-values measures the difference between the sample and population means. The proportion of variance explained by the models is presented as Marginal R<sup>2</sup> (variance explained by fixed effects alone) and Conditional R<sup>2</sup> (variance explained by both fixed and random effects). Model fit was assessed using the Akaike Information Criterion (AIC) and the Bayesian Information Criterion (BIC), where lower values indicate a better trade-off between model fit and complexity.

**Supplemental Table 4:**
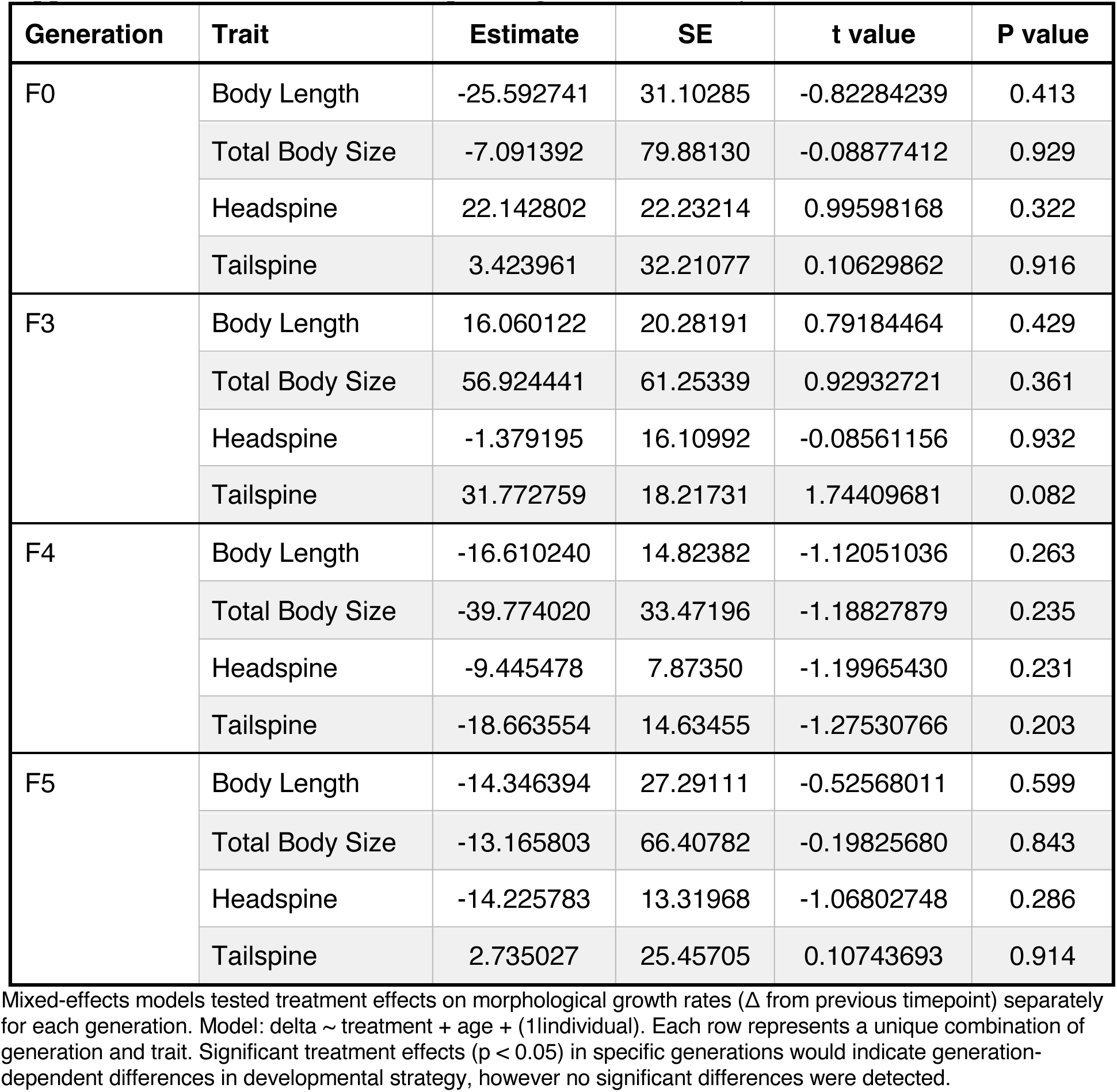
Generation-specific growth rate analysis.

**Supplemental Table 5:** Retrospective power analysis for F5 generation.

|  | Effect Sizes (µm) |  |  | F5 Sample Sizes |  | Power Analysis |  |  |  |
| --- | --- | --- | --- | --- | --- | --- | --- | --- | --- |
| Spine | F3 Effect | F4 Effect | Avg Effect (µm) | F5 n Control | F5 n Induced | Cohen's d | Power | Needed n (80%) | Est. Power |
| Head Spine | 37.0 | 43.8 | 40.4 | 8 | 41 | 0.57 | 0.30 | 49 | 29.8% |
| Tail Spine | 8.2 | 85.4 | 46.8 | 8 | 41 | 0.96 | 0.66 | 19 | 66.4% |

Effect sizes for head and tail spine traits were averaged across F3 and F4 generations and used as benchmarks to estimate statistical power for detecting similar effects in F5 given actual sample sizes (n = 8 control, n = 41 induced). Cohen's *d* represents standardized effect size (treatment effect divided by pooled SD). Columns report observed effect sizes (µm), sample sizes, calculated power, and the estimated sample size required to achieve 80% power.

